# LiverDCP: A Disease–Cell–Protein Framework for Multi-scale Modeling of Disease Biology

**DOI:** 10.64898/2026.08.07.743628

**Authors:** Ziyu Shi, Zhiyuan Song, Heather Stevenson-Lerner, Bingning Dong, Haiqing Zhao

## Abstract

Understanding how molecular interactions give rise to disease phenotypes across cellular contexts remains a central challenge in biomedical research. Here, we introduce a Disease–Cell–Protein (**DCP**) paradigm for modeling multi-scale disease biology, which jointly represents disease states, cellular composition, and protein interaction networks within a unified graph architecture. We instantiate this paradigm in the liver as **LiverDCP** by integrating LiverHomo, a harmonized single-cell atlas of liver diseases, with proteome-wide predicted protein-protein interactions to construct over 280 context-specific interactomes across diverse liver disease and cellular conditions. LiverDCP employs a multi-context representation learning strategy that enables joint training across hundreds of disease-cell environments, capturing shared interaction principles while preserving context-specific variation. LiverDCP incorporates pretrained protein sequence-derived features through a geometry-aware two-phase training scheme that preserves embedding structure while improving predictive performance. The resulting DCP protein embeddings reveal extensive rewiring of protein functional states across diseases, providing a transferable representation for downstream biomedical applications. Without GWAS supervision during representation learning, LiverDCP enables disease-risk gene classification and identifies cell types through which genetic risk may act. For therapeutic target discovery, LiverDCP recovers established Phase II+ MASH targets and prioritizes previously unrecognized candidates from the unannotated proteome, with 26 of the top 50 predictions showing independent PubMed evidence related to MASH biology. Context-specific interaction analysis further provides mechanistic hypotheses for less-characterized candidates. Together, these results establish DCP as a generalizable framework for connecting molecular interactions, cellular context, genetic risk, and therapeutic opportunities across complex diseases.

## Introduction

Protein function is not intrinsic to sequence alone but is shaped by cellular context and interaction networks^1,2^. In complex diseases such as liver disorders, including metabolic dysfunction-associated steatotic liver disease (MASH) and hepatocellular carcinoma (HCC), pathological processes emerge from coordinated interactions across diverse cell types within the hepatic microenvironment^3–5^. The liver is a central immunometabolic organ, integrating metabolic and immune functions within a tightly regulated cellular environment^6,7^. Disruption of this balance leads to chronic inflammation, fibrosis, and tumorigenesis^3,8^. Despite advances in genomic profiling^9–11^, therapeutic development for liver diseases remains challenging, in part because conventional approaches fail to account for the context-dependent nature of protein function.

Deciphering the molecular wiring of these pathologies requires moving beyond static canonical pathways to understanding protein function as an emergent property of interaction networks at cellular resolution. Early network-based studies established that protein function can be inferred from connectivity patterns within protein-protein interaction (PPI) networks^12–15^, and that disease phenotypes arise from perturbations of these networks rather than isolated gene effects^16–18^. Subsequent work further demonstrated that disease-specific interactomes—such as cancer protein interaction landscapes^19,20^—encode critical functional rewiring underlying pathogenesis. More broadly, pathway- and network-based frameworks have highlighted that biological processes are organized into interconnected modules, in which protein function is inherently context-dependent^21^.

Recent advances in single-cell RNA sequencing (scRNA-seq) have provided the requisite parts list, cataloging cellular heterogeneity across liver states^22,23^. However, translating these transcriptional atlases into actionable interactomes remains a formidable computational challenge^24–26^. While protein structural prediction tools^27–30^ such as AlphaFold and protein language models^20–24^ like ESM have revolutionized our understanding of isolated protein properties, they largely neglect the cellular environment in which proteins function. In parallel, emerging context-aware graph neural networks, such as PINNACLE^36^ and GATSBI^30^, as well as single-cell representation models^34,38,32,33^ (e.g., scVI, scGPT), have begun to capture aspects of cellular heterogeneity. Moreover, recent efforts to integrate multi-modal data have further improved protein function prediction^39,40^. Nevertheless, existing approaches remain limited in their ability to jointly model protein interaction topology, disease-state-specific network rewiring, and intrinsic protein properties within a unified framework^35^.

To overcome these limitations, we developed **LiverDCP**, an implementation of the Disease–Cell–Protein (DCP) framework that models protein function through context-aware interaction networks across diseases and cell types. As its single-cell foundation, we constructed LiverHomo, a harmonized transcriptomic atlas spanning diverse human liver diseases and cellular populations. By integrating disease- and cell-type-specific expression profiles from LiverHomo with proteome-wide structural interaction priors from PrePPI-AF, our previously developed structure-informed PPI prediction framework^43,44^, we constructed over 280 context-specific interactomes representing distinct disease–cell environments. These interactomes provide the molecular context for multi- context representation learning, enabling LiverDCP to model how protein functional states vary within the dynamic landscape of disease-specific cellular environments.

The core innovation of LiverDCP lies in its disease-informed multi-scale graph architecture, which models biological context across disease, cellular, and protein levels. Unlike prior graph-based approaches that learn representations within individual cellular contexts^45,46^, LiverDCP employs a multi-context representation learning strategy that enables joint training across hundreds of disease-cell environments, capturing shared interaction principles while preserving context-specific variation. At the architectural level, context-specific PPI networks are coupled with a disease-cell metagraph, thereby integrating molecular and cellular signals across scales. To further incorporate intrinsic protein properties, we integrate pretrained sequence-derived features through a geometry-aware two-phase training scheme that stabilizes representation learning while preserving context-specific embedding structure. Rather than focusing solely on predicting individual interactions, LiverDCP learns high-dimensional protein embeddings that encode disease- and cell-type-specific functional variation, providing a scalable framework for integrating multi-omics data and interpreting disease mechanisms.

In the Results section, we first describe the cellular landscape captured by LiverHomo and the architecture of LiverDCP. We then show that the learned DCP embeddings encode disease-associated cell-type contexts in latent space. Finally, we demonstrate the utility of these representations in downstream applications, including prioritizing high-risk Genome-Wide Association Study (GWAS^47^) genes based on inferred cell-type importance and rediscovering therapeutic targets for liver diseases.

## Results

### LiverHomo Defines Cellular Contexts across Liver Diseases

To systematically profile the cellular ecosystem of the human liver across diverse disease contexts, we constructed the LiverHomo atlas, a harmonized single-cell transcriptomic resource spanning healthy and diseased liver tissue. The atlas integrates publicly available scRNA-seq datasets encompassing normal liver, MASH, HCC, alcohol-associated liver disease (ALD), cirrhosis, primary biliary cholangitis (PBC), primary sclerosing cholangitis (PSC), and intrahepatic cholangiocarcinoma (iCCA). In total, six datasets were obtained Gene Expression Omnibus (GEO)^48^, three from Sequence Read Archive from (SRA)^49^, together with the Tabula Sapiens consortium^50^. After quality control, normalization, and harmonization, the resulting atlas comprises a unified resource of 1.16 million high-quality single cells (Fig. 1A).

**Figure 1.**
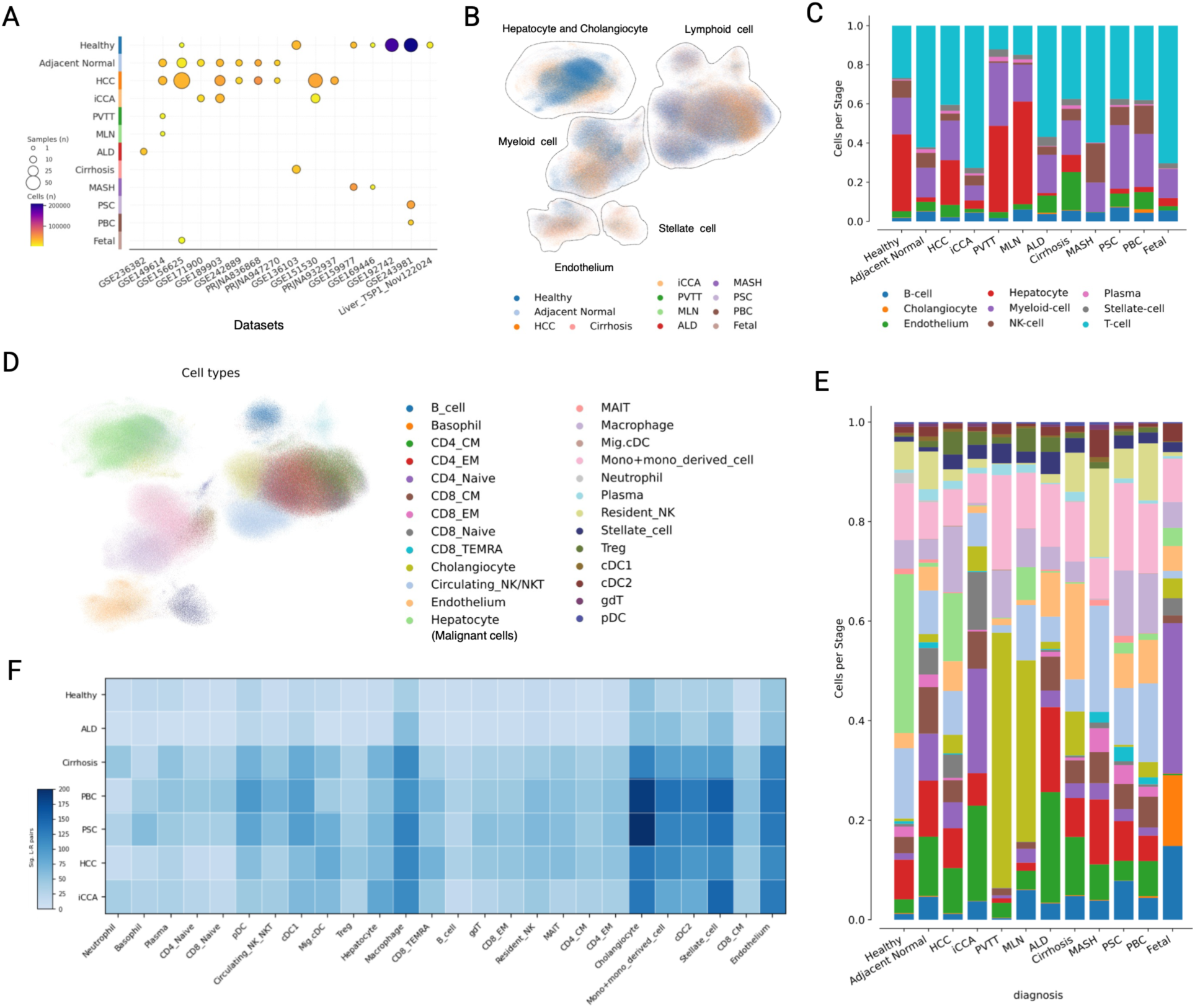
Landscape of the LiverHomo single-cell atlas for liver disease. (A) Dot plot illustrating the distribution of scRNA-seq samples from 16 public datasets (GEO/SRA) across 12 distinct hepatic conditions, spanning healthy controls, metabolic disorders (MASH, ALD), autoimmune diseases (PBC, PSC), and malignancies (HCC, iCCA). (B) Global atlas of 1.16 million harmonized single cells visualized using Uniform Manifold Approximation and Projection (UMAP), colored by clinical diagnosis. (C) Stacked bar chart displaying the relative abundance of nine major cell lineages in different diagnosis conditions. (D) High-resolution cell-type annotation visualized by UMAP, resolving fine-grained cellular heterogeneity and identifying distinct immune subsets and stromal populations. (E) Subpopulation compositional analysis, highlighting shifts in granular cell-types and revealing disease-specific expansions and contractions within immune and stromal niches. Color scheme shares with (D). (F) Altered intercellular communication shown by heatmap, quantifying the significant ligand–receptor interactions between hepatocytes and other cell-types across various clinical states.

Following Harmony-based integration, cells clustered primarily by biological identity rather than dataset of origin, while retaining variation across disease conditions, demonstrating effective batch correction (Fig. 1B). Nine major cell lineages were identified using established marker genes (Fig. S1) and automated classification via CellTypist^51^. Their relative abundance varied substantially across disease states, with prominent changes in immune and stromal populations in fibrotic and malignant liver diseases (Fig. 1C). Further subtype-level analysis revealed additional disease-associated cellular heterogeneity (Fig. 1D-E).

Beyond cellular composition, we inferred the intercellular signaling interactions integrating the cell expression profiles of LiverHomo with the curated database of known ligand–receptor pairs from CellPhoneDB^52^. This approach enabled systematic identification of potential signaling interactions between major liver cell types across disease conditions (Fig. S2). As an example, we quantified the number of predicted interactions between hepatocytes and other cellular compartments across major liver diseases (Fig. 1F). The analysis revealed a common increase in communication between hepatocytes and stellate cells, as well as cholangiocytes, suggesting enhanced signaling exchange between parenchymal and stromal populations during disease progression.

### LiverDCP: a Disease–Cell–Protein Graph Framework

#### Construction of Cell-type-specific PPI Networks

Although the LiverHomo atlas reveals the cellular composition and intercellular communication landscapes across liver diseases, understanding how disease-associated signals propagate on the molecular level requires detailed PPIs within specific cellular contexts. Recent advances in AI-based structural prediction have enabled proteome-scale inference of protein interaction networks. In our previous work, we developed PrePPI-AF^44^ and ZEPPI^43^, which integrates structural modeling and evolutionary signals to predict millions of high-confidence human PPIs, substantially expanding the coverage of experimentally derived interaction databases^53–56^. To model protein function within cellular contexts, we constructed cell-type–specific PPI networks by integrating genome-scale interaction predictions with cell-type–resolved expression profiles from LiverHomo. We first obtained a comprehensive human PPI network from PrePPI-AF, filtered it, and projected it onto cell-type–specific gene expression profiles, retaining interactions between proteins expressed in each cell type (Fig. 2A, left). This procedure generates over 286 disease-cell-specific PPI networks, enabling the representation of protein interaction landscapes across distinct cellular environments in liver diseases. These contextualized interaction networks provide the structural foundation for the LiverDCP framework described below.

**Figure 2.**
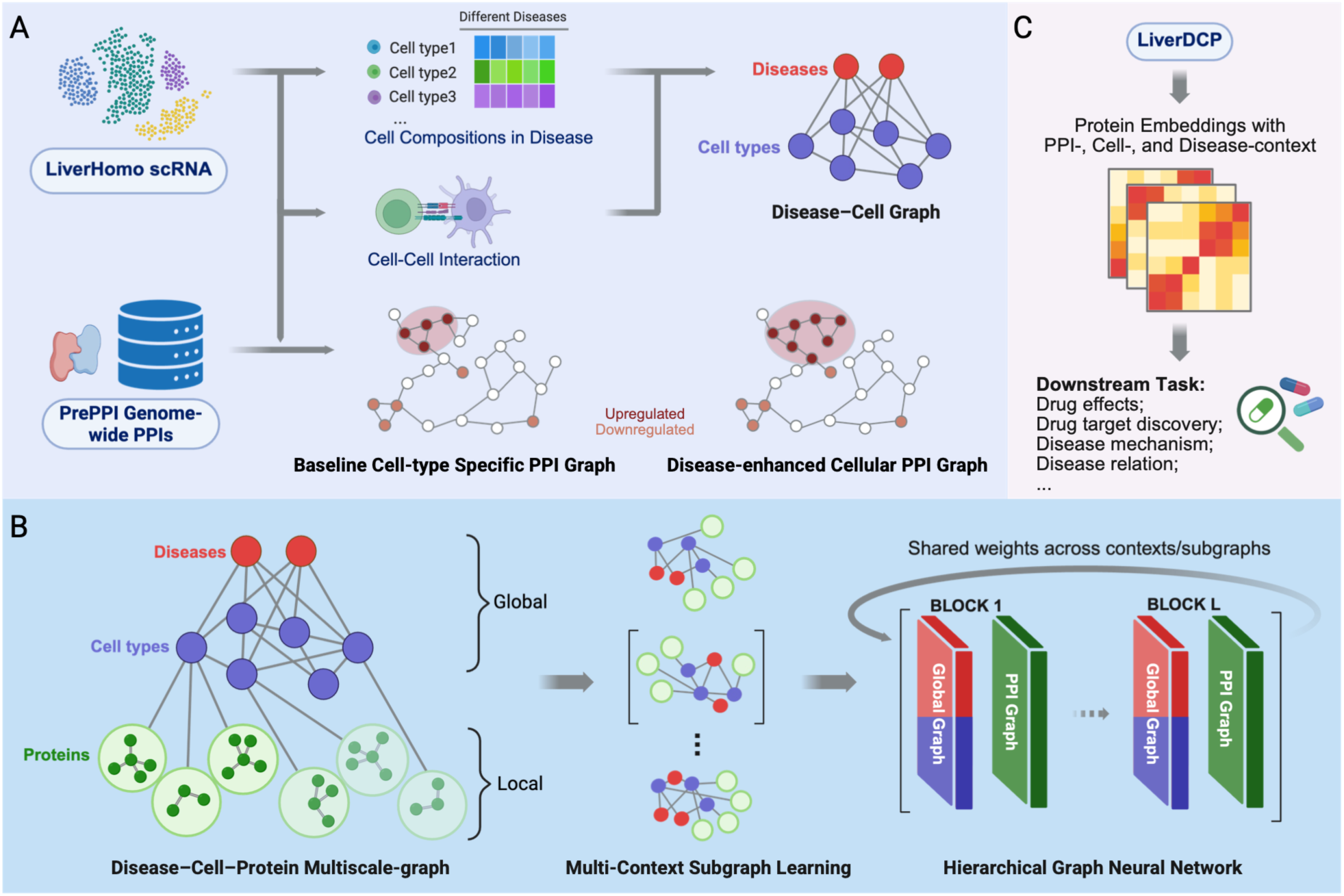
Overview of the LiverDCP multi-scale learning framework. (A) Construction of context-aware interactomes and metagraphs. Single-cell data from LiverHomo were used to quantify cell-type composition across diseases and infer cell-cell interactions via intercellular communication analysis, forming the disease-cell metagraph. In parallel, differential gene expression was used to define cell-type-specific and disease-associated gene sets, which were integrated with genome-wide structural PPI priors from PrePPI to construct context-specific PPI network graphs, including a baseline cell-type–specific PPI graph and a disease-enhanced (upregulated) PPI graph capturing pathology-driven rewiring. (B) Deep hierarchical graph learning architecture. The LiverDCP model integrates the hierarchical priors of (A) into a unified disease-cell-protein metagraph, bridging global (disease and cell-type) and local (protein) network topologies. The architecture utilizes multi-context subgraph learning driven by a hierarchical graph neural network. This network employs Global and Local modules, sharing weights across contexts and subgraphs to ensure robust, generalized feature extraction while preventing overfitting to specific niches. (C) LiverDCP generates protein embeddings inherently infused with protein interactions, cellular context, and disease states, enabling diverse downstream analyses, including drug effects, target discovery, and disease mechanism elucidation.

#### Graph Representations of Disease, Cell and Protein

The power of LiverDCP lies in its hierarchical representation of liver pathophysiology, which is captured through multi-scale graph structures: a global metagraph representing clinical and cellular relationships, and a collection of cellular PPI Networks representing molecular interactomes (Fig. 2A). The metagraph (**g**_MG_) is a directed heterogeneous graph designed to represent the relationships between disease states and cellular constituents (Fig. 2A right). It consists of disease and cell nodes that represent the diverse cell types in a specific liver condition. Features of a cell node include compositions that quantify the proportional abundance of that population within the specific disease. The connectivity within **g**_MG_ is defined by heterogeneous edge types, **ε***_MG_* , comprising **ε***_DC_* and **ε***_CC_*. We define bipartite edges **ε***_DC_* between diseases and their observed cell types, alongside disease-specific CCI edges **ε***_CC_*. At the local scale, LiverDCP deploys a protein graph **g**_k_ to represent PPI networks in a specific cell type, such as Macrophages, in a cirrhosis environment. **g**_k_ is treated as a homogeneous graph where the nodes are proteins and edges represent PPI interactions. This multiplicity of networks allows the model to capture how the interactome signature of one protein shifts depending on its cellular and clinical niche. To ensure statistical robustness, networks with fewer than 100 nodes are excluded, and all node indices are deterministically mapped to a global protein universe to enable consistent feature retrieval.

#### Model Architecture and Learning Strategy

LiverDCP employs a hierarchical Graph Neural Network (HGNN)^57^ architecture that jointly models protein interactions across cellular and molecular scales through a unified Disease-Cell-Protein graph (Fig. 2B). The DCP framework operates on two coupled components: a global disease–cell metagraph ( **g**_MG_ ) that encode relationships among disease states and cellular environments, and context-specific PPI networks (**g_k_**) that capture local interaction topology.

In **g**_MG_, disease and cell-type nodes are initialized using cell-type proportion data derived from single-cell deconvolution. This ensures that the starting state of the metagraph reflects the actual cellular composition of the diseased liver tissue. In **g**_k_, PPI networks are highly context dependent, with interaction patterns varying across cell types and disease conditions. To capture this heterogeneity, we initiate the protein node features from a Gaussian-like random distribution to provide a baseline for structural learning, allowing the model to learn DCP embeddings driven primarily by the topology of context-specific interactomes. Node representations emerge from connectivity patterns rather than fixed biological priors (e.g., protein sequences), enabling the model to focus on learning the disease-cell-specific interaction landscapes.

At the core of the DCP model framework is a shared graph attention network (GAT^58^) backbone coupled with a semantic attention mechanism for multi-scale context integration. The model performs iterative information propagation between context-specific PPI networks and the disease–cell metagraph, enabling effective graph fusion and cross-scale representation learning (Fig. 2B). Across all PPI contexts, the GAT layers share a common set of attention parameters to enable consistent modeling of interaction patterns, while context-specific multilayer perceptron (MLP) projection heads preserve biological heterogeneity across disease–cell conditions. During the global phase, semantic attention within the Graph Fusion Convolution (GFC) module^59^ aggregates updated disease and cell node representations within the metagraph, allowing the model to capture higher-order relationships across biological contexts. In the subsequent local phase, the refined metagraph embeddings are propagated to protein nodes within each interactome, enriching local protein representations with disease– and cell-type–specific contextual information. By coupling local interaction topology with higher-order biological organization, this hierarchical propagation framework enables the model to learn context-aware protein embeddings that reflect both molecular interaction structure and cellular environment.

To capture the diversity and relatedness of protein functions across heterogeneous disease states, LiverDCP introduces a multi-context representation learning strategy that enables joint representation learning across diverse disease–cell environments. Unlike prior frameworks such as PINNACLE that train models independently within individual cellular contexts, LiverDCP simultaneously learns from a controlled collection of context-specific PPI networks which are coupled through a shared subgraph and unified model parameters (Fig. 2B). This design allows the model to capture both context-specific interaction patterns and shared molecular principles across related biological states. As a result, LiverDCP learns protein representations that generalize across contexts while retaining disease– and cell-type–specific interaction structure. This controlled transfer enhances robustness to data sparsity and enables knowledge sharing across related biological states, providing a foundation for downstream analyses in heterogeneous systems.

### Disease-specific Protein Representations in DCP Embeddings

A central output of LiverDCP is a set of protein embeddings that encode disease– and cell-type–specific interaction context within a unified representational space. By learning multi-context representations from context-dependent variations in protein interaction networks, the model captures how protein function is reshaped across cellular environments and disease states, providing a foundation for downstream analyses such as target discovery and mechanistic interpretation (Fig. 2C).

To assess the latent structure of the learned representation space, we examined whether these DCP embeddings reflect the functional plasticity of the liver proteome by analyzing the geometry of its latent space. LiverDCP was trained until convergence, as indicated by stabilization of validation AUROC, and embeddings from the final epoch were used for downstream analysis (Fig. S3). In total, the model generated 777,609 context-specific protein representations (12,085 unique proteins) spanning 286 unique disease-cell context niches (11 diseases, 26 cell-types).

The learned DCP embedding space exhibits clear organization by cellular and disease context, with well-separated embedding clusters corresponding to distinct biological contexts. For example, hepatocytes from HCC and Healthy tissues form distinct, non-overlapping clusters despite sharing much of the underlying proteome (Fig. 3A). This separation indicates that the model has learned to encode the functional interactome, where a protein like ALDOB or PCK1 (marked in Fig. 3A networks) occupies a different geometric position depending on whether it is functioning in a metabolic, fibrotic, or neoplastic context. Topologically distinct PPI subnetworks (extracted top 20 nodes) confirm that the local neighborhoods of key metabolic enzymes are fundamentally rewired in disease states. Within a single disease type, the overall DCP embeddings across different cell types exhibit topological diversity and connections that undergo context-dependent pattern shifts compared with other disease conditions (Fig. S4).

**Figure 3.**
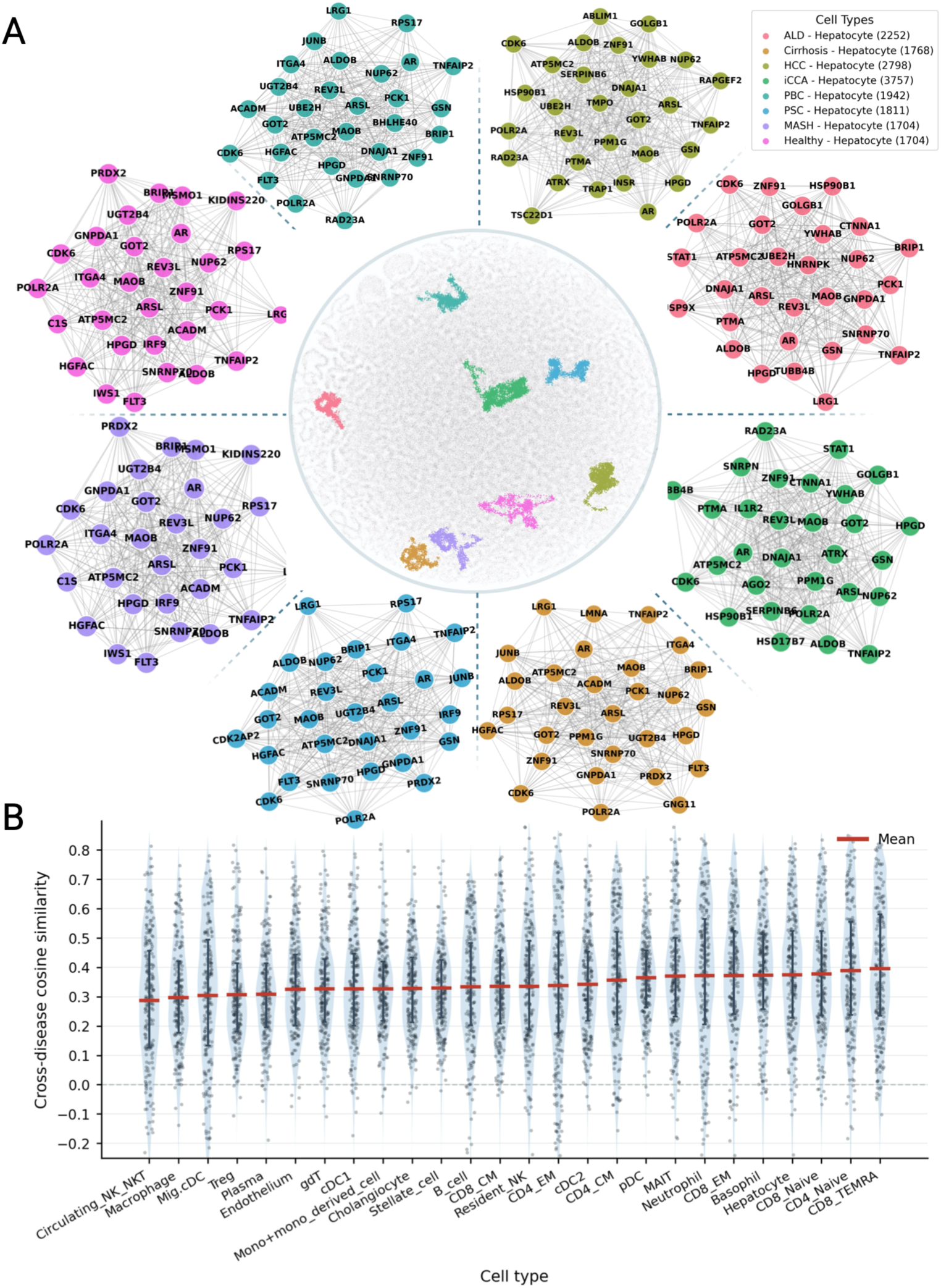
Resolving disease-specific protein function via context-aware embeddings. (A) Geometric separation of hepatocyte functional states. UMAP visualization of hepatocyte protein embeddings across multiple liver conditions (e.g., healthy, MASH, HCC) shows distinct clustering in latent space despite shared protein identities, indicating that the model captures disease-specific functional interactomes. Corresponding PPI networks of top high-degree proteins illustrate local neighborhood rewiring of key metabolic enzymes in disease states. (B) Quantification of protein functional plasticity across diseases. Box plots show the distribution of cosine similarity for proteins within each cell type, where each point represents the cross-disease similarity of an individual protein (computed across pairwise disease comparisons). Values near 1 indicate stability whereas near 0 indicate divergence.

To further detail this functional plasticity, for a given protein in a given cell-type, we calculated the cosine similarity by comparing its embedding from one disease to that from another. Similarity values were averaged across pairwise disease comparisons and summarized at the cell-type level (Fig. 3B). Cell types were ranked by mean similarity in ascending order, with lower similarity indicating greater context-dependent embedding divergence. to identify the most disease-rewired populations. Immune populations undergoing pathological remodeling, such as circulating NK/NKT cells, exhibit the highest embedding divergence (lowest similarity), reflecting rapid transitions in functional state and validating that LiverDCP learns dynamic, context-specific representations rather than static features.

Across all cell types, the overall cross-disease cosine similarities of all proteins span approximately from −0.2 to 0.85, while the mean cross-disease similarity per cell-type ranges from ∼0.29 (circulating NK/NKT) to ∼0.40 (CD8 TEMRA). Taking endothelium cells as an example (mean = 0.32; Fig. 3B, Table S1), their housekeeping proteins such as HNRNPK, UQCR10, and TMSB10, retain markedly high cross-disease similarity (0.531, 0.533, and 0.543, respectively), reflecting their constitutive structural and metabolic roles. In contrast, context-dependent signaling proteins such as LYVE1, EGR1, and CRIM1, have shown substantially low similarity (0.151, 0.136, and 0.127, respectively), consistent with their disease-specific remodeling and transcriptional reprogramming across liver pathologies. Specifically, cell-types deeply involved in pathological remodeling and dynamic functional state transitions, including circulating NK/NKT cells, macrophages, migratory cDCs, regulatory T cells, plasma cells, and endothelium, exhibit the lowest mean cross-disease similarity (0.286–0.325; left side of Fig. 3B), reflecting extensive context-specific rewiring during inflammation and immune response. In contrast, relatively quiescent, naive or terminally differentiated populations, including CD8 TEMRA, CD4 naive, and CD8 naive T-cells, together with hepatocytes, retain higher similarity across disease contexts (0.374–0.395; right side of Fig. 3B), consistent with more stable or transcriptionally constrained baseline states. This quantitative gradient validates that LiverDCP does not learn a static canonical embedding for each protein, but rather a dynamic, context-aware representation that adapts to the specific molecular wiring of the disease microenvironment.

### Sequence-Augmented Protein Representations

The DCP framework is designed to learn protein interaction topology and embed cell-type-specific PPI networks into protein representations, enabling their direct use in downstream tasks with medical and therapeutic relevance. From a complementary perspective, protein function is also shaped by intrinsic biochemical and evolutionary constraints encoded in amino acid sequences. Recent advances in protein language models, such as ESM^32^, have demonstrated that large-scale sequence pretraining can encode rich biochemical and structural knowledge within protein embeddings. These representations capture evolutionary constraints and functional motifs that are largely invariant across cellular contexts, making them a natural complement to topology-driven interactome modeling. To incorporate this complementary source of information, we augmented protein nodes in LiverDCP with embeddings derived from a pretrained ESM model.

However, directly initializing node features with ESM embeddings leads to a collapse of context-specific structure, as identical sequence representations across cell types reduce the model’s ability to learn cellular context-specific distinctions, resulting in degraded predictive performance (AUROC ∼0.50). To address this limitation, we introduced an adapter-based integration strategy together with a two-phase geometry-aware training procedure that enables the gradual incorporation of sequence-derived information into topology-driven representations (Fig. 4A). During Phase I, the graph encoder and zero-initialized Progressive Sequence Adapter (PSA) are jointly optimized to establish stable context-specific representations while monitoring embedding quality. Once the embedding threshold is reached, the graph encoder is frozen and the same PSA is further reinforced and optimized in Phase II, allowing protein sequence information to refine the learned representations without disrupting the topology-driven embedding geometry (see Methods for details).

**Figure 4.**
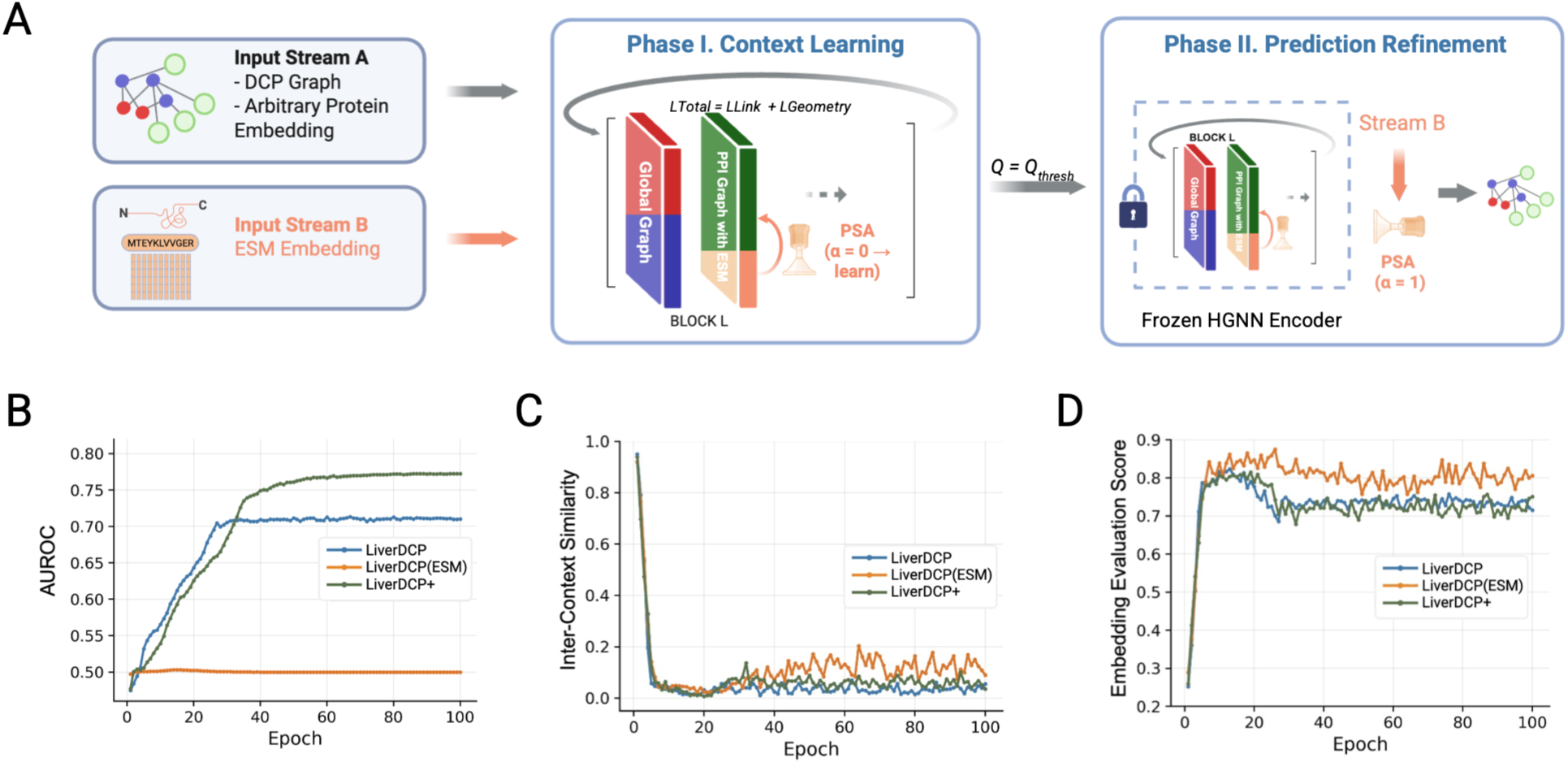
Sequence integration through progressive sequence adapter preserves context-specific embedding geometry while improving predictive performance. (A) Overview of DCP and protein sequence integration via two-phase geometry-aware learning strategy. In Phase I, the graph encoder and PSA are jointly trained. Once the embedding quality threshold is reached, the graph encoder is frozen and the same PSA is further optimized to incorporate protein sequence information for improved link prediction while preserving context-specific representations. The resulting sequence-augmented model is referred as LiverDCP+. (B) Link prediction performance (validation AUROC). LiverDCP+ (green) achieves superior predictive accuracy (ROC-AUC ∼0.77), significantly outperforming the topology-only baseline LiveDCP (blue, ∼0.71) and the sequence-initialized model LiveDCP(ESM) (orange, ∼0.50). (C) Embedding evaluation score dynamics. The hybrid architecture (green) keeps the high clustering quality of the topological baseline (blue) while integrating sequence data, whereas the ESM-only initialization (orange) yields suboptimal embedding geometry. (D) Inter-context differentiation. LiverDCP+ rapidly minimizes inter-context cosine similarity (green), effectively disentangling distinct cellular states with the same efficiency as the topological baseline (blue), while the sequence-only model (orange) struggles to differentiate protein functions across diverse cell types.

To systematically evaluate this integration, we assessed model performance using not only AUROC (Fig. 4B) but also with two complementary metrics that characterize the learned representations: an embedding evaluation score (cluster coherence) and inter-context cosine similarity (context separability) (Fig. 4C-D). We compare three model variants: the native topology-driven model (LiverDCP), a sequence-initialized baseline (LiverDCP(ESM)), and the sequence-integrated model (LiverDCP+). LiverDCP+ achieves the highest AUROC (0.77), outperforming both LiverDCP and LiverDCP(ESM) (0.71 and 0.50), while maintaining comparable embedding evaluation score and inter-context specificity (Fig. 4C-D, S5). Together, these results indicate that combining interaction topology with sequence-derived information yields protein representations that are both context-specific and biologically informative. Subsequent analyses are based on this sequence-integrated model unless otherwise specified.

### DCP Embeddings Enable Context-Specific Interpretation of GWAS Risk

GWAS have cataloged thousands of genetic variants associated with human diseases, most of which reside in non-coding regions and require mapping to candidate genes. However, even after gene-level assignment, translating these loci into mechanistic insights remains challenging due to limited tissue- and cell-type–specific context. To address this limitation, we evaluated whether LiverDCP embeddings improve the prioritization of disease-relevant genes. Specifically, we assessed the capacity of LiverDCP embeddings to distinguish high-confidence risk genes and target proteins from housekeeping controls across seven major liver pathologies, including MASH, ALD, Cirrhosis, HCC, iCCA, PBC, and PSC. To isolate the specific contribution of the liver-specific graph topology, we benchmarked performance against protein sequence language model ESM^32^.

#### Superiority of Context-Aware Representations

Across all seven disease indications, models incorporating liver-specific graph priors (LiverDCP and LiverDCP+) consistently outperformed sequence-only baselines (ESM). In MASH, a complex metabolic pathology, the sequence-integrated LiverDCP+ achieved an AUROC of 0.946, substantially higher than that of the ESM baseline (0.790). Similarly, in ALD, the topology-driven LiverDCP model achieved strong discrimination (AUROC 0.970), whereas the ESM model performed significantly worse (AUROC 0.776) (Fig. 5C, S6; Table S2). These results indicate that incorporating cell-type–specific interaction context provides substantial gains in predicting disease-associated genes beyond sequence-derived representations alone.

**Figure 5.**
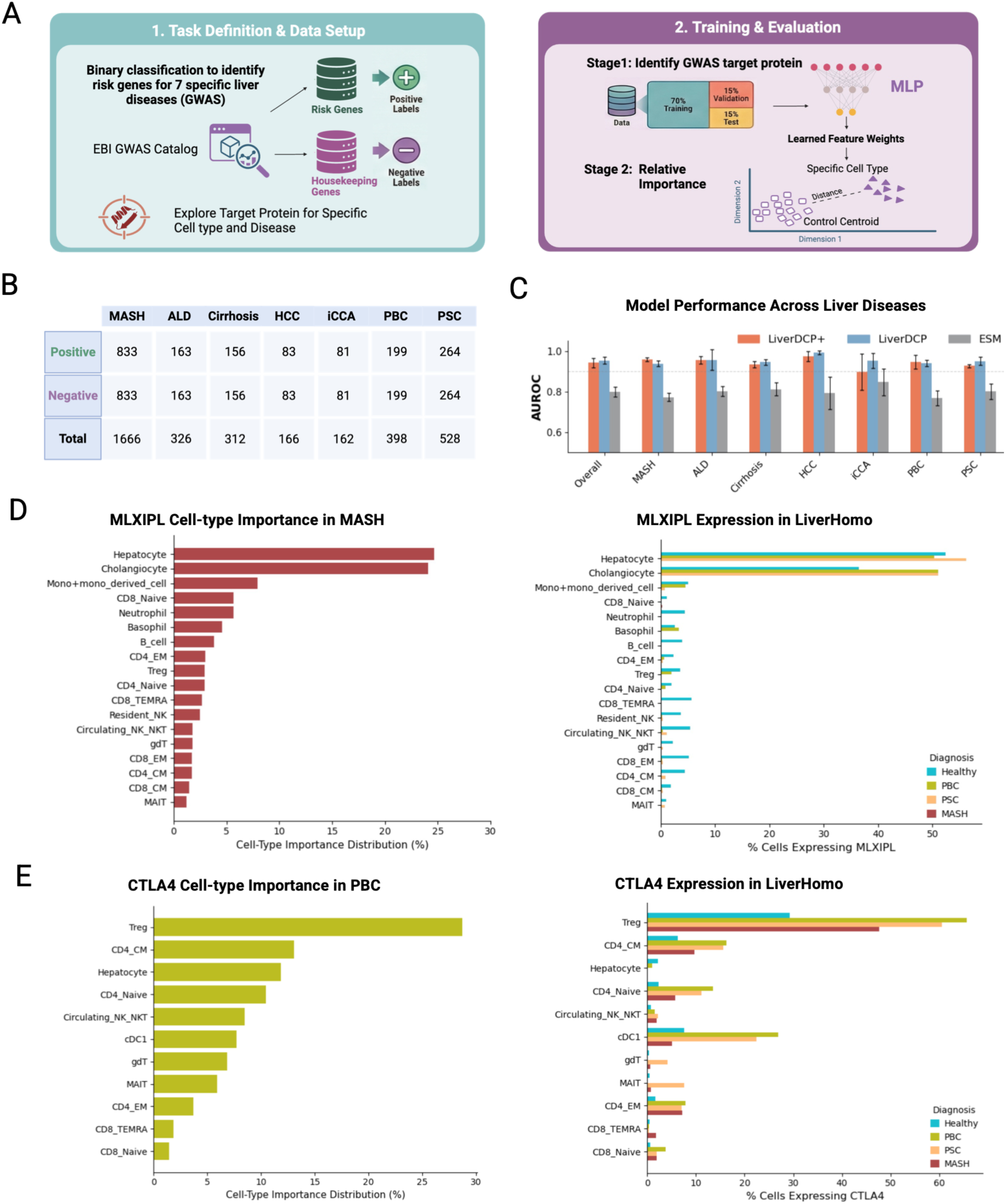
LiverDCP+ Decodes Disease-Specific Risk Genes via Contextualized Embeddings. (A) GWAS fine-tuning pipeline. Schematic of the predictive framework, including data setup for the binary classification of risk versus housekeeping genes (left) and the two-stage training strategy for identifying target proteins and cell-type importance (right). (B) Dataset composition. Counts of positive (disease-associated) and negative (housekeeping) target genes were evaluated across seven liver pathologies. (C) Model performance. AUROC comparison demonstrating that graph-aware models (LiverDCP+, orange; LiverDCP, blue) consistently outperform the sequence-only baseline (ESM, grey) across all diseases. Error bars represent standard deviation (n=5 seeds). (D- E) Cell-type importance and expression for validated risk loci. Predicted cell-type importance distributions (left) and corresponding scRNA expression frequencies (right) for (D) the metabolic risk gene MLXIPL in MASH, prioritizing hepatocytes and cholangiocytes, and (E) the immune checkpoint gene CTLA4 in PBC, prioritizing regulatory T-cells.

Unlike protein language models, which encode intrinsic sequence features, LiverDCP embeddings capture context-dependent functional variation by modeling protein interactions within cellular environments. As a result, the model distinguishes how the same protein functions across distinct cell types, reflecting the underlying wiring of disease pathways. The strong performance of both LiverDCP and LiverDCP+ indicates that the primary signal for genetic risk is encoded in interaction topology. While predictive performance between the topology-driven and sequence-integrated variants is comparable, we use the sequence-integrated embeddings for downstream analyses to leverage their incorporation of intrinsic protein features.

#### Cell-type Importance for MLXIPL in MASH

Using sequence-integrated LiverDCP embeddings, MLXIPL (encoding ChREBP) is identified as a highly ranked disease-associated gene in MASH. Consistent with its role as the liver’s primary glucose sensor and driver of *de novo* lipogenesis (DNL)^60,61^, the model prioritized hepatocytes as the primary cell type (Fig. 5F). This is consistent with the established role of ChREBP as a key regulator of hepatic glucose metabolism and de novo lipogenesis (DNL), processes that contribute to lipid accumulation during steatosis^60,62^.

Beyond this established mechanism, the model further assigns high MLXIPL importance to cholangiocytes, suggesting a role in the metabolic demands of the ductular reaction. This association likely captures the metabolic demands of the “ductular reaction”, the liver’s regenerative response to chronic injury^63^. Reactive cholangiocytes function analogously to cancer cells, undergoing a metabolic switch to aerobic glycolysis (the Warburg effect) to support rapid proliferation^64^. The upregulation of MLXIPL in this context is a functional necessity, providing the transcriptional drive for the glycolytic flux required for ductular expansion^65^. Together, these results demonstrate that LiverDCP links genetic signals to cell-type–specific metabolic programs, capturing both established and context-dependent disease mechanisms.

#### Cell-type Importance for CTLA4 in PBC

In PBC, using sequence-integrated LiverDCP embeddings, the immune checkpoint CTLA4 is identified as a top disease-associated target. Reflecting its role as a master regulator of T cell activation and immunotolerance, the model prioritized regulatory T cells (Tregs) as the dominant cellular context (∼30% importance, Fig. 5E). This is supported by LiverHomo, which shows robust, disease-specific high CTLA4 expression in PBC Tregs. Within the immune compartment, the model also captures the broader immunological network by prioritizing circulating NK/NKT cells and central memory CD4+ T cells. The capturing of CD4_CM reflects the trans-endocytosis mechanism where Treg-expressed CTLA4 strips CD80/CD86 from antigen-presenting cells to dampen effector responses^66,67^. These findings link genetic susceptibility at the CTLA4 locus to specific immune cell populations and regulatory mechanisms, illustrating how LiverDCP resolves global genetic risk into localized cellular circuits driving disease pathology.

### DCP Embeddings Enable Therapeutic Target Discovery

Having shown that LiverDCP embeddings provide mechanistic insights into disease-associated genes, we next evaluated their translational utility for drug target discovery. We curated a high-confidence pharmacological atlas spanning seven major liver diseases by integrating data from the OpenTargets Platform^68^, restricting to proteins targeted by therapeutic agents that reached Phase II or later clinical trials. This procedure yielded 488 unique clinical targets. As a comparison, we constructed a 1:1 matched set of proteins that are chemically druggable (defined as having at least one drug in clinical trials in the OpenTargets Platform) but are not documented Phase II+ clinical targets for any of the seven liver diseases studied and are not GWAS risk genes for these conditions^36^.

We next established a unified validation and target prioritization framework based on the geometric organization of known targets in the DCP embedding space (Fig. 6A). Using 5-fold cross-validation, we treated 10% of the known targets in each fold as held-out positives and used the remaining 90% to define the disease-specific target distribution. For each held-out target, we computed its Mahalanobis distance to the embedding distribution of the remaining known targets, thereby quantifying how closely it aligned with the disease-target cluster. To benchmark the performance, we then did the same for 506 non-target proteins.

**Figure 6.**
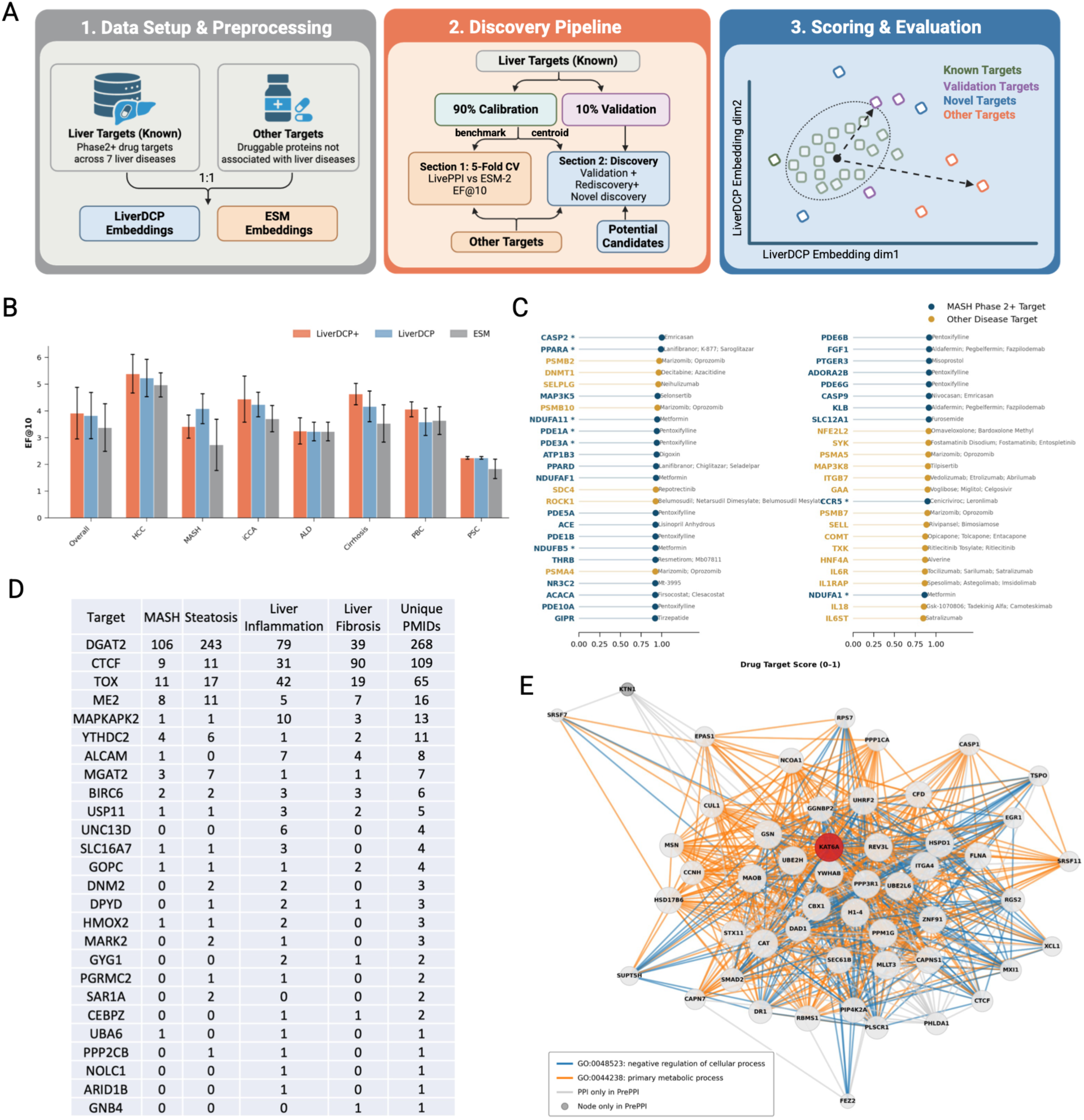
Unified validation pipeline and performance of LiverDCP in liver disease therapeutic target discovery. (A) Pipeline for the drug target discovery encompasses embedding generation (LiverDCP vs. ESM), a 90:10 stratified split for cross-validation, and candidate ranking via Mahalanobis distance to the known target centroid. (B) Cross-validation performance. Graph-aware models (LiverDCP+ in orange, LiverDCP in blue) achieve superior early target enrichment (EF@10) compared to the sequence-only baseline (ESM in grey) across seven liver diseases (mean ± std., n=5 folds). (C) Target rediscovery in MASH. Mahalanobis scoring successfully prioritizes the clinically approved MASH target THRB, alongside top-ranked inflammatory targets (MAP3K8, IL6ST) with repurposing potential. (D) PubMed evidence for the top 50 novel targets prioritized by LiverDCP was systematically assessed across MASH, fibrosis, inflammation, and steatosis. (E) PPP2CB-centered protein interaction network within macrophages. Edge colors represent distinct GO biological processes, highlighting PPP2CB’s localized coordination with mediators of lipid-driven inflammation and selective autophagy (e.g., ALOX5AP, OPTN), as well as established macrophage stress and surface regulators (e.g., ACTN1, ENTPD1).

Candidate proteins were ranked by this distance-based score, and performance was assessed using the enrichment factor (EF), which quantifies the concentration of held-out true targets among the top-ranked predictions relative to random expectation. We benchmarked LiverDCP+ against context-agnostic ESM-2 sequence embeddings under the same evaluation scheme. Across the seven liver diseases, LiverDCP+ demonstrated consistently stronger early enrichment than the sequence-only baseline, outperforming ESM-2 in 6 of 7 indications as measured by the EF at rank 10 (EF@10) (Fig. 6B, S6). For example, in MASH, LiverDCP+ achieved an EF@10 of 3.93 (Table S3), compared with 2.38 for ESM-2, indicating that the topological structure of tissue-specific interactions captures critical pharmacological liabilities beyond sequence information alone.

### Drug Target Rediscovery in MASH

Moving beyond validation, we evaluated the model’s ability to prioritize high-confidence, clinically actionable candidates for drug-target rediscovery in MASH (Fig. 6C, S7). Notably, the model ranked THRB (Thyroid Hormone Receptor Beta) at rank 25. THRB is the molecular target of resmetirom^69^, the first and currently only FDA-approved pharmacotherapy for MASH, demonstrating that LiverDCP+ can recover established therapeutic targets from a large, unannotated proteomic space. This result is consistent with the central role of THRB in regulating hepatic lipid metabolism and mitigating lipotoxicity^70^. Among the top 50 prioritized candidates, the model also identified several targets more commonly associated with other disease indications, including MAP3K8 and IL6ST, both of which are involved in inflammatory signaling pathways^71,72^. These findings suggest that LiverDCP+ can highlight biologically relevant targets and facilitate the identification of potential drug-repurposing opportunities through shared metabolic and inflammatory mechanisms.

### Novel Target Discovery in MASH

We next expanded the MASH-trained LiverDCP model across 7268 of unannotated proteomes to identify previously unrecognized therapeutic candidates. For the top 50 proteins prioritized by LiverDCP+ (Table S4), we then systematically examined PubMed literature evidence to evaluate their liver biological relevance, focusing on MASH and its three major processes: steatosis, inflammation, fibrosis (Fig. 6D). Among these candidates, 26 had prior literature evidence in at least one category, spanning a broad spectrum from extensively studied proteins to largely unexplored candidates. DGAT2 showed the strongest support, with 268 publications across the four categories, followed by CTCF (109), TOX (65), and ME2 (16). Several additional candidates, including MAPKAPK2, YTHDC2, ALCAM, and MGAT2, showed more limited but relevant evidence, whereas many predictions had little or no prior connection to MASH-associated biology. These results indicate that LiverDCP can recover biologically supported candidates beyond established Phase II+ liver therapeutic targets while simultaneously identifying underexplored proteins that may provide new hypotheses for therapeutic investigation.

### Contextual Characterization of PPP2CB in MASH Macrophages

Among the top-ranked novel targets was the catalytic subunit of protein phosphatase 2A, PPP2CB (Table S4), a signaling regulator with mechanisms distinct from canonical kinase-centered pathways. In contrast to several highly ranked candidates with substantial prior literature support (Fig. 6D), PPP2CB remains largely unexplored in MASH, with limited evidence linking it to MASH-associated processes. We therefore investigated whether LiverDCP could provide cellular and functional context for this less-characterized candidate by examining PPP2CB-centered interaction networks across MASH-associated cell types in LiverHomo, followed by Gene Ontology Biological Process (GO:BP) enrichment analysis (Fig. 6E).

It turns out that this is the most cohesive, disease-relevant signaling sub-network that emerged specifically within a macrophage-centric inflammatory context, a key immune population driving lipotoxicity and fibrogenesis in MASH^4^. Within this framework, PPP2CB forms a dense interactome coordinating mediators of lipid-driven inflammation and selective autophagy (ALOX5AP, OPTN)^74,75^ with established macrophage regulators (ACTN1, ENTPD1)^76,77^. Gene Ontology (GO) enrichment validated this functional specialization, highlighting significant associations with response to stress (GO:0006950), cell death (GO:0008219), and hemopoiesis (GO:0030097). These architectural features suggest PPP2CB acts as a key regulatory node constraining inflammatory cascades in activated macrophages, providing a promising role as the drug target for MASH. Together, these network features suggest that PPP2CB may function as a regulatory hub within inflammatory macrophage programs in MASH, supporting its potential as a therapeutic target. More broadly, these results illustrate LiverDCP’s ability to place broadly expressed proteins into disease-relevant cellular interaction contexts, enabling mechanistically interpretable target discovery.

## Discussion

Protein functions in complex diseases are inherently context-dependent, yet most computational approaches model proteins as context-independent entities defined primarily by sequence or static structure^45,78^. In this study, we present LiverDCP as a unified DCP framework that models protein function as an emergent property of disease- and cell-specific interaction networks. By integrating protein interaction topology with cellular context derived from single-cell transcriptomics, LiverDCP enables the representation of proteins within a Disease-Cell-Protein space that captures both local protein-protein connectivity and global pathological environments. This multi-scale formulation allows the model to resolve functional variation of proteins across disease states, moving beyond gene-centric or bulk-level interpretations. As a result, LiverDCP provides a principled approach for linking molecular interactions to cellular phenotypes, offering a new lens through which protein function can be interpreted in complex tissue systems.

A central component enabling the LiverDCP framework is the LiverHomo atlas, which provides a harmonized single-cell view of liver disease across diverse pathological states. While prior scRNA-seq studies have revealed substantial cellular heterogeneity in individual liver conditions, they are typically analyzed in isolation and lack integration with PPI interaction networks^22,23,79,80^. By systematically integrating over one million cells across multiple liver diseases, LiverHomo enables consistent characterization of cellular composition and transcriptional programs within a unified reference space. Importantly, coupling these data with structural protein interaction priors allows the reconstruction of cell-type-specific interaction networks that reflect disease-associated rewiring of the hepatic microenvironment. In this way, LiverHomo serves not only as a comprehensive resource of liver cellular states, but also as a foundation for linking gene expression to protein interaction topology in a disease- and cell-type-resolved manner.

Beyond methodological advances, LiverDCP enables biologically interpretable insights into disease-associated protein function across cellular contexts. For instance, our model identifies MLXIPL (ChREBP) in MASH as a top-ranked risk gene and assigns the highest importance to hepatocytes, consistent with its established role in hepatic glucose sensing and *de novo* lipogenesis^60,62,81^. Notably, LiverDCP also highlights cholangiocytes as a secondary context, suggesting that MLXIPL-associated metabolic programs may extend beyond parenchymal hepatocytes, particularly during ductular reaction and tissue remodeling^61^. In immune-driven pathology, the model prioritizes PPP2CB within macrophages, a core population driving lipotoxicity and fibrogenesis^82^, where its interaction network links lipid-driven inflammation and selective autophagy with the regulation of cellular stress^75,76^. These examples demonstrate that LiverDCP not only recapitulates established biology but also uncovers context-specific associations that remain obscured in bulk or transcriptome-only analyses.

At the translational level, LiverDCP demonstrates strong potential in disease-gene interpretation and therapeutic target discovery. By embedding proteins within specific disease- and cell-type interaction contexts, the model improves the classification of GWAS-derived risk genes, addressing a central limitation of genetic studies in which functional interpretation is often decoupled from cellular context^46^ (Fig. 5). Crucially, LiverDCP assigns cell-type-specific importance to risk genes, providing a direct link between genetic association and cellular mechanism. Beyond genetic interpretation, our target discovery analyses demonstrate how these context-aware representations can systematically prioritize therapeutically relevant proteins. LiverDCP successfully rediscovered established Phase II+ MASH targets (Fig. 6C) and, when extended to 7,268 proteins without such therapeutic annotations, identified a distinct set of previously unrecognized candidates (Fig. 6D). Systematic literature analysis of the top 50 predictions revealed that many already have independent evidence linking them to MASH or major pathological processes including fibrosis, inflammation, and steatosis, despite not being established Phase II+ liver therapeutic targets (Fig. 6E). Importantly, the predictions also included substantially less-characterized proteins, illustrating the ability of LiverDCP to move beyond existing therapeutic knowledge and generate new hypotheses. PPP2CB provides one such example: although it has little prior literature linking it to MASH-associated processes, its context-specific interaction network places it within a macrophage-associated inflammatory program, providing a potential cellular and molecular basis for further investigation. Thus, rather than prioritizing proteins in isolation, LiverDCP connects target ranking with the disease, cell type, and molecular network in which a candidate may exert its pathological function. More broadly, the learned DCP embeddings provide a transferable representation that can be leveraged to predict potential therapeutic effects and guide target selection across related disease contexts^83^. Together, these results position context-aware protein embeddings as a unifying interface for connecting genetic variation, cellular mechanisms, and therapeutic intervention in complex diseases.

The performance of LiverDCP highlights the importance of integrating complementary sources of biological information within a unified representation. Protein interaction topology provides a relational framework that captures how proteins function within cellular networks, while single-cell transcriptomic context defines the cellular environments in which these interactions occur. Sequence-derived embeddings from protein language models further encode intrinsic biochemical and evolutionary constraints that are largely invariant across conditions. Each modality offers a partial view of protein function; their integration yields a more complete representation that reflects both molecular identity and contextual regulation. Crucially, naïve incorporation of sequence features can obscure context-specific structure, as identical sequence embeddings across cell types reduce the model’s ability to distinguish cellular environments. To address this, LiverDCP employs a controlled two-phase learning strategy through PSA that can progressively incorporate sequence-derived information into topology-driven representations, preserving cell-type specificity while enhancing representational expressiveness. This multi-scale design—combining topology, cellular context, and sequence priors—provides a principled DCP framework for modeling protein function in complex, heterogeneous biological systems.

Despite these advances, several limitations remain. First, LiverDCP relies on single-cell transcriptomic data and the PPIs derived from PrePPI-AF, both of which have inherent constraints. Single-cell RNA-seq datasets remain subject to technical noise, dropout effects, and incomplete detection of lowly expressed genes. While PrePPI-AF provides broad proteome-wide coverage, the underlying interactions are computationally inferred rather than directly measured in specific cellular contexts. As a result, inaccuracies in the interaction scaffold may propagate into downstream representations. Improving interaction accuracy and incorporating experimentally resolved, cell-type-specific interactomes—potentially through emerging multi-omics modalities such as spatial transcriptomics and single-cell proteomics—will be important future directions^84^. Second, the current DCP framework focuses primarily on genes upregulated in diseased cells, thereby enhancing signal-to-noise ratio but potentially overlooking functionally important downregulated genes or loss-of-function mechanisms. Third, although single-cell transcriptomic data provide valuable context, gene expression does not always reflect protein abundance or activity due to post-transcriptional and post-translational regulation^85,86^. Large-scale proteogenomic studies have shown that while mRNA-protein correlations are broadly positive^19,87–89^, their magnitude is typically moderate and varies substantially across biological processes^90^. For example, metabolic and immune pathways tend to exhibit higher concordance, whereas processes such as ribosome biogenesis, RNA splicing, and oxidative phosphorylation show weaker correspondence between transcript and protein levels^89^. As a result, transcriptome-derived context may incompletely capture the quantitative and functional state of proteins, particularly for pathways under strong post-transcriptional control^90,91^. Integrating complementary modalities, such as single-cell proteomics and spatially resolved measurements, will be important for improving the biological fidelity of context-specific interaction modeling. Finally, while LiverDCP embeddings show strong performance in downstream tasks, experimental validation remains necessary to confirm newly predicted disease associations and therapeutic targets. Addressing these limitations by integrating orthogonal data types, including spatial transcriptomics, proteomics, and longitudinal datasets, will further enhance the accuracy and interpretability of context-aware protein modeling.

In summary, LiverDCP provides a unified DCP framework for modeling protein function as a context-dependent property shaped by cellular environment and interaction networks. Integrating single-cell transcriptomics with protein interaction topology and sequence-derived information enables systematic protein–cell alignment mapping of disease-associated proteins within specific cellular contexts. Importantly, LiverDCP functions as a context-aware protein representation encoder, generating embeddings that capture both intrinsic biochemical properties and disease-and cell-type–specific interaction environments. These representations are inherently transferable across downstream tasks, offering a generalizable interface for protein-centered applications, including disease gene identification, drug target prioritization, and prediction of therapeutic effects. Beyond liver disease, this DCP framework is readily extensible to other complex tissues where cellular heterogeneity and network rewiring are central. As multi-omics datasets continue to expand, incorporating spatial, temporal, and proteomic dimensions will further enhance the resolution of context-aware protein modeling, positioning LiverDCP as a scalable foundation for linking molecular interactions to cellular phenotypes in precision medicine.

## Method

### LiverHomo Single-Cell Atlas

#### Acquisition and Preprocessing of scRNA-seq Data

To construct a harmonized single-cell atlas of liver disease, we developed LiverHomo by integrating publicly available scRNA-seq datasets across diverse pathological conditions. Human liver scRNA-seq datasets were systematically curated from public repositories, including: (i) the NCBI Gene Expression Omnibus (GEO)^48^; (ii) the Sequence Read Archive (SRA); and (iii) the Tabula Sapiens consortium^50^. Raw sequencing data were processed using a standardized pipeline. FASTQ files were aligned to the human reference genome (GRCh38)^92^ with CellRanger (v7.1.0)^93^, generating Gene expression count matrices for further analysis. This unified preprocessing workflow ensured comparability across all datasets.

#### Quality Control, Integration, and Cell-type Annotation

Gene expression matrices were analyzed using Scanpy from the scverse framework^94^. Standard quality control procedures were applied to remove low-quality cells^95^ and potential doublets^96^. Data were normalized, log-transformed, and highly variable genes were selected for downstream analysis. Batch effects across datasets were corrected using Harmony^97^, followed by dimension reduction^98^ and Leiden clustering^99^. Cell types were annotated using a hybrid strategy combining automated label transfer using the CellTypist^51^ with cluster-specific differentially expressed genes (DEGs) analysis^95^ and canonical lineage markers compiled from literature and the CellMarker database^100–106^. Selected populations were further resolved at the subtype level, including T-cell annotation using starCAT^107^. Detailed quality-control criteria, integration parameters, and annotation procedures are provided in the Supplementary Methods.

#### Inference of Cell-Cell Communication Networks

To characterize intercellular communication, ligand–receptor interaction was inferred using CellPhoneDB^52^. Candidate interactions were defined based on curated ligand–receptor pairs and filtered according to expression within the corresponding cell populations. Statistical significance of CCI was assessed using permutation testing, combined with repeated subsampling to ensure robustness. Consensus CCI networks were constructed for each disease condition by retaining interactions consistently observed across iterations and subsequently incorporated into the cellular context represented by LiverDCP.

### Construction of Context-Specific Protein Interactomes

To construct cell-type– and disease-resolved PPI networks, we first derived two complementary gene sets from the LiverHomo atlas. Disease-associated DEG genes were identified for each condition relative to healthy controls using the memento-de method, while cell-type–specific marker genes were identified by contrasting the expression profiles of each cell type with those of all other cell types in the healthy cohort.

These gene sets were then mapped onto a proteome-scale interaction prior generated by PrePPI-AF, a structure-informed PPI database that integrates AlphaFold-predicted monomer structures with non-structural evidence to compute likelihood ratios (LRs) for PPIs^44^. This approach enables genome-wide PPI coverage with reduced experimental context bias compared to curated or high-throughput interaction datasets (e.g., HINT^108^, STRING^109^, BioGRID^110^, HuRI^55^), which may be influenced by assay-specific or literature-driven biases. PrePPI-AF has previously reported over 1.3 million high-confidence human PPIs. In this study, interactions over an LR cutoff of 200 were retained as the prior network, leading to 1.9 million PPIs.

For each cell type, two complementary PPI subnetworks were constructed by intersecting the prior network with the gene sets defined above: (i) a baseline subnetwork comprising interactions among cell-type–specific marker genes, representing constitutive cellular wiring, and (ii) disease-specific subnetworks comprising interactions among genes upregulated in each disease condition, capturing pathology-driven rewiring. Interactions were retained only among genes with detectable expression in the corresponding cell-type and were further prioritized by PrePPI-AF LR score and structural compatibility

### LiverDCP Model

#### Construction of the Heterogeneous Metagraph and Homogeneous PPI-graph

The metagraph is defined as a directed heterogeneous graph **g***_MG_* = (**V***_MG_*, {**ε***_m_*}_m∈{CC,DC}_), where nodes **V***_MG_* include disease-type nodes (**V***_D_*) and cell-type nodes (**V**_C_), and edges ({**ε***_m_*}_m∈{CC,DC}_) include disease-cell associations (**ε***_DC_*) and intercellular communication (**ε***_CC_*) derived from the LiverHomo atlas. Edges **ε***_DC_* were constructed based on cellular composition within each disease, retaining cell types with at least 10 cells and excluding adjacent normal samples, with edge weights normalized by relative abundance. Edge **ε***_CC_* were inferred from disease-specific consensus ligand–receptor networks and consolidated across conditions; interactions shared by multiple diseases were annotated accordingly and weighted by their frequency of occurrence. Pairwise Jaccard similarity between disease-specific interaction networks was further computed to quantify the extent of shared versus distinct intercellular communication programs. In parallel, each global context (a specific disease-cell combination) is associated with a local PPI network, represented as a homogeneous graph **G***_k_*= (**V***_k_*, **ε***_k_*) capturing the interaction topology among expressed proteins.

#### Model Architecture

The DCP model framework is designed to model protein function within disease- and cell-type–specific contexts. The model operates on a hierarchical representation that integrates two complementary graph structures: (i) a global metagraph capturing relationships among diseases and cell types, and (ii) a local network graph of context-specific PPIs representing molecular interactions within each disease**-**cell pair. This design is conceptually related to recent context-aware graph frameworks such as PINNACLE, which model protein interactions across tissue and cellular hierarchies, but extends these approaches by explicitly incorporating disease context and integrating multi-scale biological information with protein sequence-derived features.

The model architecture consists of *L* stacked graph fusion blocks that coordinate representation learning across two topological scales: the global disease-cell metagraph **g***_MG_* and the local intracellular PPI networks **g**_k_. At block *l,* the module updates the previous hidden representations of the global metagraph 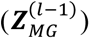 and the local context-specific protein interactomes 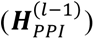 through two coupled global and local propagation modules, formalized as:

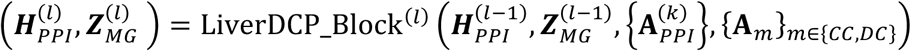

The global module serves a dual purpose: (1) it aggregates multi-relational context within the disease–cell metagraph; and (2) pulls local PPI graph signals upward. In the metagraph branch, the model processes the diverse relations—specifically cell-cell (**ε**_CC_ ) and disease-cell (**ε**_DC_ ) edges—using a relation-wise GAT kernel. To weigh the importance of these different biological relations, the per-relation outputs are fused using a semantic attention mechanism (SemAtt**_β_**). This process occurs in two successive stages using shared semantic attention parameters but independent GAT kernel:

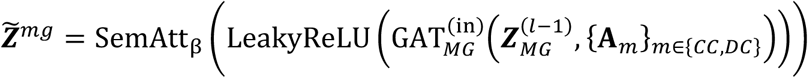

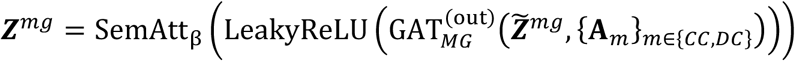

In parallel, the PPI branch operates independently for each unique (disease, cell-type) context *k*. The protein features are convolved via a shared GAT to capture robust interactome motifs. Crucially, these features are then transformed by a context-specific, two-layer multi-layer perceptron (MLP^(k)^ ), which projects the shared topological learning into a unique biological embedding subspace for that specific cellular environment to yield the intermediate state ***H***^(k)^.

The local module consumes the outputs of the global module to refine the context-specific protein representations. For each context *k*, the local module integrates three complementary components: (1) self-update (***G***^(k)^), consisting of a dedicated GAT convolution over the local PPI network to capture immediate interactome topology; (2) top-down contextual contribution (***m***^(k)^), wherein metagraph-derived context embeddings are broadcast back down to the protein level through mean-pooling over the specific contextual neighborhood 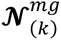 in metagraph; and (3) residual connection, a linear projection (via **W**) of the input representation to preserve original signals and prevent oversmoothing. The refined PPI representation is defined as:

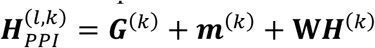

This architecture enables that the final protein embeddings are dynamically shaped by both their immediate local molecular interactions and the higher-level clinical context of the disease-cell metagraph. Full mathematical formulation and implementation details are provided in Supplementary Information.

#### Multi-context Representation Learning

LiverDCP adopts a multi-context representation-learning framework in which a shared graph backbone is optimized jointly across diverse disease-cell interactomes. Here, each context is defined as a disease-cell pair and training is performed on stochastic groups of 50 contexts and their context-specific PPI networks. During training, all contexts in one context group are jointly propagated through the graph fusion blocks with conventional mini-batching (size=64). Iterative exposure to diverse disease-cell environments enables LiverDCP to jointly learn shared interaction patterns while preserving context-specific variation.

#### Training Objective

LiverDCP is trained using a knowledge-guided link prediction objective derived from provided edge priors. Following graph encoding, node embeddings are decoded through relation-specific scoring function to estimate interaction logits and probabilities for binary cross-entropy (BCE) optimization:

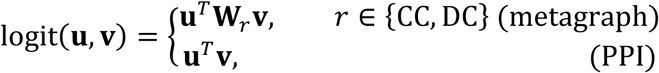

where **u** and **v** denote node embeddings, **W***_r_* is a per-relation learnable weight matrix applied elementwise. For PPI interactions, logits are computed through dot-product similarity between protein embeddings. Model parameters are optimized using a joint link prediction objective across both PPI networks and disease-cell metagraph. The overall loss is defined as: ℒ_link_ = *θ* · ℒ_pp1_ + (1 − *θ*) · ℒ_MG_, where θ is a tunable parameter. ℒ_ink_ and ℒ_MG_ denote the BCE functions for protein-level and metagraph-level interactions, respectively. This joint objective encourages the model to learn representations that are coherent across molecular and cellular scales, while preserving higher-order disease and cell-type structure. To promote generalization, edges are partitioned into training, validation, and test sets, and complementary edge masking is applied during training to prevent information leakage between message passing and supervision.

#### Hierarchical Edge Partitioning and Complementary Masking

To ensure robust generalization, we implement a two-tiered edge splitting protocol for both PPI interactomes **g***_k_* and the metagraph **g**_MG_. At the dataset level, PPI edges are independently partitioned within each context using an 80:10:10 train/validation/test split, whereas metagraph edges are partitioned globally using an 85:5:10 ratio. To address class imbalance in the sparse PPI networks, dynamic negative sampling is applied during training by uniformly sampling non-interacting protein pairs within the same context at a 1:1 ratio relative to positive edges. During each training iteration, the available training edges are further divided using a complementary masking strategy. Specifically, 50% of the training edges are designated as message-passing edges defining the GNN graph topology, while the remaining 50% are withheld as supervision targets for link prediction. This prevents trivial self-prediction by ensuring that the model is prohibited from seeing the exact interactions it is tasked to infer during the convolutional forward pass.

### Embedding Evaluation Metrics

To assess whether LiverDCP embeddings capture biologically meaningful and context-specific representations, we evaluated the embedding geometry using complementary metrics that measure disease-specific rewiring, inter-context separability, and cluster coherence.

#### Cross-disease Protein Similarity

To quantify functional rewiring across disease contexts, we performed cross-disease protein similarity analysis within each cell type. For a given cell-type *c* in *k* disease states {*d*_1_, … , *d*_k_} ( *k* ≥ 2 ), we enumerated all 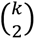 unordered disease pairs. For each disease pair (*d*_A_, *d*_D_) , we identified the set of shared proteins 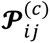 and sampled up to 500 proteins for computational efficiency. For every sampled protein *p*, we computed the cosine similarity between its disease-specific embeddings: 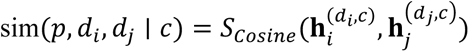 where 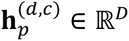 denotes the embedding vector of protein *p* in disease *d* and cell type *c*. Per-cell-type aggregate statistics were derived as: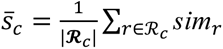, where **R**_c_ is the set of all pairwise similarity records for cell type *c*.

#### Inter-context Cosine Similarity

To evaluate separability between context-specific embeddings, we computed the mean pairwise cosine similarity between context centroids. For each context *k*, the centroid embedding is defined as: 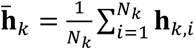This metric *Scosine* is defined as: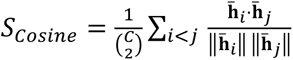 , where *S_cosine_* near zero or negative indicate strong context separation, while values approaching 1 suggest embedding collapse.

#### Silhouette Score

Cluster compactness and separation were further evaluated using the silhouette coefficient. For each protein embedding *x*_i_with context label ℓ_i_, the silhouette coefficient is 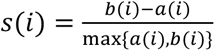 where *a*(*i*) is the mean intra-cluster distance and *b*(*i*) is the mean distance to the nearest foreign cluster. The overall silhouette score 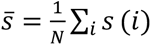 ranges from −1 to 1 , with higher values indicating denser, better-separated clusters.

#### Composite Embedding Evaluation Score

To jointly evaluate embedding separation and cluster coherence, we defined a composite embedding evaluation score *Q* ∈ [0,1]:

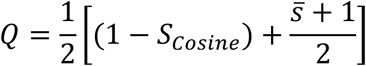

where the first term captures centroid-level separation and the second normalizes the silhouette score to [0,1]. In ESM adapter, this score served as the primary signal for triggering the Phase I/II transition: when *Q* exceeded a threshold *Q*_thresh_ (default 0.7), the best-quality checkpoint was saved. The quality score was also monitored alongside AUROC to detect regressions in cluster structure during link-prediction optimization.

### Infusing Pretrained Protein Sequence Feature

To complement topology-driven learning with intrinsic protein features, we incorporated sequence-derived representations from the pretrained ESM-2 model^32^. However, naïve integration of pretrained sequence embeddings can conflict with context-specific, topology-driven representations, as sequence features primarily encode conserved, context-free information. To address this, we designed a zero-initialized adapter (PSA) that introduces sequence information in a controlled manner. Specifically, PSA projects the fixed 480-dimensional ESM-2 embeddings ( **e**^ESM^) through a low-rank bottleneck with a learnable gate, initialized to ensure the ESM contribution is null at the beginning. This design allows the model to first establish stable topology-driven representations and subsequently incorporate evolutionary information in a gradual and data-adaptive manner via a learnable scale parameter *α* , enabling the model to dynamically balance structural and evolutionary signals. For protein *i* in the *k*^th^ disease-cell context, the fused input feature 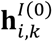 is:

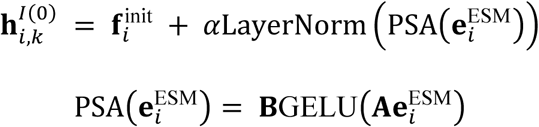

where **A** ∈ ℝ^r x 480^ (*r* = 48) is a down-projection matrix (Kaiming initialized), **B** ∈ ℝ^d x r^ (*d* = 1024) is an up-projection matrix strictly initialized to zero, and 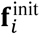 denotes Gaussian-like random distribution.

### Two-phase Training Strategy

We observed a mismatch in optimization timescales: context-specific embedding emerges early in training, whereas continued optimization toward the link prediction progressively degrades the learned embedding space, reducing separability across different cellular contexts (Fig. S8). To resolve this, we employed a two-phase geometry-aware training strategy. Phase I focuses on learning topology-driven representations while gradually incorporating sequence features through the ESM PSA via adjusting the scaling parameter α.

In Phase I (context learning), we introduced two complementary geometric constraints, ℒ_center_ and ℒ_diversity_, balancing the topological reconstruction (ℒ_link_) and geometric regularization. The total loss ℒ_total_ is then defined as a weighted sum of three components:

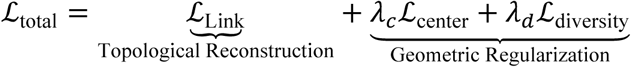

The center loss ℒ_center_ is the intra-class compactness. Defined as 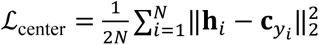 we penalize the Euclidean distance between a protein’s embedding **h**_A_ and its context-specific centroid **c**_yi_, forcing the model to cluster functionally related proteins tightly within their cellular manifold. The diversity regulation term ℒ_diversity_describes the inter-context separability. To ensure that different cell types maintain distinct representations, we maximize the orthogonality between context centroids. The diversity loss penalizes high cosine similarity between the centroids of different contexts **c** and 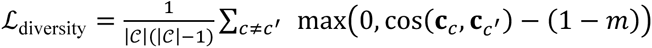, where m is a margin hyperparameter that enforces a minimum angular separation.

In Phase II (link refinement), rather than freezing the encoder at a fixed epoch, the system monitors the composite embedding evaluation score *Q* (defined in the session below) throughout training, triggering the phase change dynamically. Specifically, Phase II is activated when two conditions are jointly satisfied: (i) the quality has previously peaked above the threshold by a margin of at least 0.05, confirming that the encoder has learned discriminative context structure, and (ii) the current quality has subsequently dropped to or below the threshold (*Q* ≤ *Q*_thresh_), indicating the onset of embedding degradation. After moving to phase II, the GNN encoder and its implemented PSA from phase I are frozen to preserve the learned embedding space. The PSA adapter architecture is reused, with an independent parameter set, to provide a bypass pathway for integrating ESM embeddings. A Residual Network (ResNet) is activated to operate on the frozen embedding from phase I:

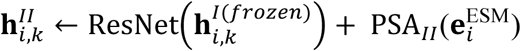

Where ResNet(**u**) = **u** + **D** · PReLU(LayerNorm(**C** · **u**)), **C**, **D** ∈ R^d×d^
Training then focuses on refining predictive performance through a new PPI link prediction head while maintaining the established balance between topology-driven and sequence-derived representations.

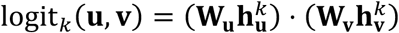

Where **u**, **v** denote node embeddings, **W_u_** and **W_v_** are two learnable projection matrices to reduce the rank of the PPI embeddings. This design enables continued optimization of predictive performance through ℒ_Link_ backpropagation without altering the underlying representation geometry. Meanwhile, the geometric regularization terms, *λ*_E_ℒ_center_ + *λ*_I_ℒ_diversity_, are still utilizing to the frozen embeddings 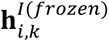.

### Training Protocol and Hyperparameters

The model was trained using the Adam optimizer and cosine annealing warm restart scheduler^111^ with a base learning rate of *η* = 1 × 10^2p^, weight decay of 5 × 10^2b^, *T*_warmup_ = 15 epochs. Training was performed on a single NVIDIA GTX 5090 GPU using PyTorch Geometric^112^. To handle the massive adjacency matrices of liver-scale interactomes (> 10^q^ edges), we implemented a memory-efficient chunked inference algorithm for the bilinear link predictor, processing edges in batches of 4,096 to constrain peak memory usage.

### Downstream Task: GWAS Risk Genes

#### Data Collection and Curation

To construct a high-fidelity ground truth dataset for liver disease risk, we integrated data from the EBI GWAS Catalog across seven major liver disease indications: MASH, ALD, Cirrhosis, HCC, iCCA, PBC, and PSC. We programmatically queried the EBI GWAS REST API for associations with *P* < 1 × 10^2q^, encompassing both genome-wide significant and suggestive loci, utilizing exact ontology matching (e.g., EFO_0004268 for PSC) and fuzzy string matching for synonymous traits. For indications with sparse GWAS representation (iCCA, PSC, Cirrhosis), we augmented the dataset using the Open Targets GraphQL API, requiring a minimum evidence score of > 0.01 derived from genetic association or somatic mutation data. All raw gene lists were harmonized to the HGNC nomenclature to filter out non-coding artifacts and resolve ambiguous loci, and genes were further filtered by requiring their presence in the liver PPI network, yielding 67–341 risk genes per disease. Finally, to mitigate class imbalance, we constructed a balanced negative set (1:1 ratio) for each disease using housekeeping genes, selected based on transcriptional ubiquity (expressed in ≥ 80 of cell types) and strict exclusion from risk lists across all seven target diseases.

#### Risk Gene Classification via MLP (Stage 1)

We formulated disease gene identification as a binary classification task. The model accepts a multi-scale input tensor combining sequence-based embeddings (ESM-2, dimension *d* = 480) and context-aware graph embeddings (LiverDCP+, dimension *d* = 256) spanning 286 cell types.

To aggregate the heterogeneous cellular contexts, we obtained the max value from each row of the embeddings for the input tensor for following MLP. Specifically, for a given gene *g* , the representation ℎ_7_ is derived by pooling the maximum activation across all *C* = 286 cell types, ensuring that a strong signal in a single relevant cell type is not diluted by non-expressing contexts:

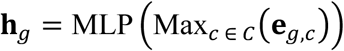

The data was split into training, validation, and test sets at the ratio of 0.70:0.15:0.15. The classifier was trained on the training set using a weighted BCE loss. The validation set was used to automatically determine when to terminate model training. The results shown in Figure 5 were from the evaluation of the trained model using the test set.

#### Cell-type Deconvolution via Weighted Geometric Distance (Stage 2)

While Stage 1 identifies what genes are risky, Stage 2 resolves where (in which cell type) these risks manifest. We developed a Weighted Z-Score Distance metric that leverages the learned feature weights from Stage 1 to measure the deviation of a risk gene from the homeostatic baseline. In detail, there are four below steps.

1. Feature Importance Extraction. We extracted the weights **W** ∈ *R*^H×D^ from the first layer of the trained Stage 1 MLP, where *D* = 256 is the embedding dimension. The global importance of each embedding dimension *i* was calculated as the mean absolute weight magnitude, normalized to a probability distribution: 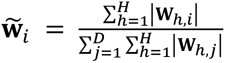
2. Control Centroid Definition. For every disease-relevant cell type *c*, we defined a homeostatic centroid based on the distribution of control (housekeeping) genes **G_ctrl_**. We computed the mean **μ**_C_ and standard deviation **σ**_C_ for each dimension: 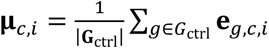
3. Weighted Z-Score Calculation. The disease relevance of a risk gene *r* in cell type *c* is defined as its weighted Mahalanobis-like distance from the control centroid. Crucially, this distance is weighted by **w**¶, ensuring that deviation is only penalized in dimensions that the Stage 1 classifier found predictive of disease risk: 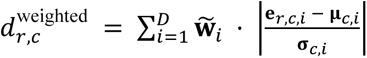
4. Relative Importance Scoring. To enable cross-cell-type comparison, weighted distances were converted into a relative importance score using a temperature-scaled exponential normalization function with a temperature parameter *τ* = 3, highlighting the specific cellular niche where the protein’s functional embedding is most perturbed: 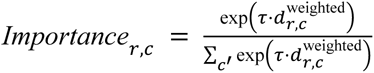

### Downstream Task: Therapeutic Target Discovery

#### Data Collection and Curation

To rigorously evaluate the translational utility of LiverDCP+, we constructed a high-confidence pharmacological atlas spanning seven major liver disease indications: HCC, MASH, iCCA, ALD, Cirrhosis, PBC, and PSC. We defined Liver Targets Labels (Known Targets) by integrating data from the OpenTargets Platform, strictly selecting proteins targeted by therapeutic agents that have reached Phase II or later clinical trials or beyond for the specific indication. This ensures the ground truth reflects clinically validated mechanisms rather than purely experimental associations. Other Disease Targets Labels (Non-Targets) were defined as proteins that are chemically “druggable” (targeted by at least one agent in OpenTargets Platform) but have no documented association with the liver disease of interest. This curated dataset yielded 488 unique clinical targets across the seven indications, providing a robust benchmark for distinguishing true therapeutic signals from the background proteome.

#### Target Representation and Mahalanobis Scoring

For each disease, protein representations were generated by aggregating cell-type specific embeddings into a unified protein vector using L2-normalized summation: 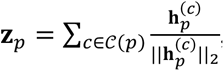 where C(*p*) denotes all the cell-types that have protein 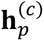 denotes the embedding of protein *p* within cell type *c*.

Candidate proteins were scored using the Mahalanobis distance relative to the distribution of known drug targets. To account for the high dimensionality of the embedding space relative to the limited number of clinical targets, we employed Oracle Approximating Shrinkage (OAS)^113^ to estimate the covariance matrix **Σ**^-1^ . The scoring function *s*(*p*) is then defined as: 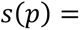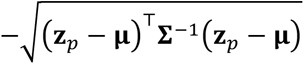 , where *μ* is the centroid of known target embeddings: 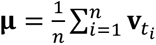 with **v**_ti_ denoting the pooled embedding of the *i*-th known target.

To obtain an interpretable ranking metric, raw Mahalanobis scores *S(p)* were normalized into a drug target score *S*_drug_ ∈ [0,1]:

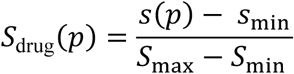

#### Internal Validation via Cross-Validated Enrichment

To quantify the specific contribution of tissue-aware topology over sequence alone, we benchmarked LiverDCP+ against the context-agnostic ESM-2 baseline using a rigorous 5-fold cross-validation (CV) framework. We partitioned the known targets into a 90% training set and a 10% held-out test set. Within the 90% training partition, we performed 5-fold CV to compute the Enrichment Factor (EF), measuring the concentration of true positives in the top-ranked predictions relative to random chance. Specifically, for the broader early-retrieval metric EF@10, this metric is defined as:

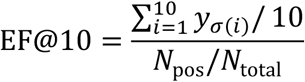

where *y_σ(i)_* indicates the ground truth label of the protein at rank *i* (sorted by *S*_drug_*(p)*), and *N*_pos_ is the total number of known drug targets in the validation fold. External Validation via Target Rediscovery

To simulate a real-world drug discovery campaign, we assessed the model’s ability to rediscover hidden targets in a large pool of novel candidates. We trained the Mahalanobis scoring model on the 90% training partition and then ranked a discovery pool comprising the 10% held-out test targets, mixed with all “novel” (negative) candidates. This setup strictly prevents information leakage, as the test targets were never seen by the covariance estimator.

We evaluated rediscovery performance using the one-sided Mann-Whitney U test to determine whether the held-out test targets received statistically higher Mahalanobis proximity scores than the novel candidates (H₁: test scores > novel scores), reporting diseases with p < 0.05 as significant. In addition, we computed the rediscovery count, the number of held-out test targets that appeared among the top-30 ranked proteins in the entire discovery pool. This metric serves as a stringent proxy for early retrieval in a prospective screening scenario, verifying that the model can prioritize clinically relevant targets above background noise.

#### PubMed Literature Evidence Analysis

To assess independent literature support for novel therapeutic candidates prioritized by LiverDCP, we systematically queried PubMed for the top 50 ranked proteins using the NCBI Entrez Programming Utilities (E-utilities). For each gene, separate searches were performed for four MASH-relevant themes: MASH, liver fibrosis, liver inflammation, and steatosis. Closely relevant theme terms with alternative names were included as keywords. Searches were restricted to gene and disease/process terms appearing in the title or abstract to reduce nonspecific matches. The number of PubMed hits returned for each gene–theme combination was recorded, and candidates with at least one matching publication were included in the literature evidence analysis (Fig. 6D). Total unique PMIDs for each target gene were counted to characterize the extent of existing literature evidence rather than as a direct measure of therapeutic relevance or experimental validation.

#### Functional Context of Target-centered Interaction Networks

To characterize the biological functional context of the target protein’s context-specific PPI network, we performed Gene Ontology Biological Process (GO:BP)^73^ enrichment analysis on all genes within the ego subgraph using g:Profiler. Enrichment significance was evaluated using Benjamini–Hochberg false discovery rate (FDR) correction with a significance threshold of 0.1 and a minimum intersection size of two genes per term. Enriched pathways were ranked by *p*-value and the top-ranked pathways were projected onto the network through colored edges. Specifically, an edge was assigned to a pathway when both endpoint nodes belonged to the corresponding enriched gene set, thereby enabling spatial localization of pathway activity within the interaction topology. Centered on the target protein, the integrated presentation of disease-enhanced PPIs, upregulated genes, and GO pathway enrichment analysis highlights the central principle of DCP paradigm: linking protein function to context-specific molecular interaction networks to enable mechanistically informed downstream explorations such as target discovery.

## Data availability

The datasets analyzed in this study and the source data underlying all main and supplementary figures have been deposited in Zenodo and are available at https://zenodo.org/records/20431227.

## Code availability

The LiverDCP source code, including codes for data preprocessing, model training, evaluation, and downstream analyses, is publicly available at https://github.com/zhaolabutmb/LiverDCP. Pretrained model weights are available through Huggingface at https://huggingface.co/zhaolabutmb/LiverDCP.

## Acknowledgements

We thank Surendra Negi for insightful discussions on model training, Xiaoqin Wu (UTHealth Houston) for suggesting GWAS resource, Omar Saldarriaga and Esteban Arroyave Sierra for help in LiverHomo database. H.Z. acknowledges the generous support of the Welch Foundation (H-2308-20260402) and the UT System Rising STARs Award.

## Author contributions

Z.S and H.Z. designed research; Z.S., Zh.S. and H.Z. performed research; Z.S., Zh.S., H.S., B.D. and H.Z. analyzed data; Z.S., Zh.S., and H.Z. wrote the manuscript; all authors revised the manuscript.

