## Supplemental Material for "LiverDCP: A Disease–Cell–Protein Framework for Multi-scale Modeling of Disease Biology" (from another version of this preprint: version 1, 2026-08-10)

### Supplementary Information

Table of Contents

Supplementary Method Details — 1

Supplementary Figures — 7

Supplementary Tables — 14

References — 17

#### Supplementary Method Details

##### LiverHomo Database Construction

*Quality Control, Dimensionality Reduction, and Annotation*

Processed gene expression count matrices were analyzed using Scanpy (v1.9.6) from the scverse framework, implemented in Python (v3.10) [1]. Quality control (QC) filtering was applied to remove low-quality cells as follows: cells with fewer than 500 or more than 6,000 detected genes were excluded, along with those exhibiting mitochondrial gene content exceeding 20% (pct_counts_mt > 20), ribosomal gene content below 3% (pct_counts_rp < 3), or hemoglobin gene content above 1% (pct_counts_hb > 1) [2]. Doublets were detected and removed using Scrublet (v0.2.3) with default parameters [3].

Following QC, the AnnData object was transferred to GPU-accelerated computation using the rapids-single cell package (GPU-accelerated via RAPIDS cuML, CUDA 12.0)on an NVIDIA RTX 5090 GPU for enhanced performance [4]. Data were normalized to a target sum of 10,000 counts per cell (target_sum=1e4) and log-transformed. Highly variable genes were selected using the following thresholds: minimum mean expression of 0.0125, maximum mean of 3, and minimum dispersion of 0.5.

Batch effects and unwanted sources of variation (e.g., mitochondrial and ribosomal gene percentages) were regressed out, followed by data scaling to unit variance. Principal component analysis (PCA) was performed retaining the top 50 components (n_pcs=50). Batch correction was applied using Harmony to integrate datasets [5]. Nearest neighbors were computed with n_neighbors=10 and the ivfflat algorithm. Dimensionality-reduced embeddings were generated via uniform manifold approximation and projection (UMAP) with a minimum distance of 1 (min_dist=1) [6]. Finally, cell clustering was conducted using the Leiden algorithm at a resolution of 0.7 [7]. T cells were subsequently isolated and further subtyped using starCAT with predefined gene expression programs[8].

Cell type annotation was conducted utilizing a hybrid approach combining automated label transfer with manual, evidence-based curation. Initially, automated cell typing was performed using the CellTypist algorithm, referencing the 'Healthy_Human_Liver' model to establish baseline transcriptomic identities[9]. To validate and refine these predictions, unsupervised clustering was performed, and cluster-specific differentially expressed genes (DEGs) were subsequently identified utilizing the 'deg_analysis' function from the omicverse package (v1.5.4)[2].

To confirm cluster identities, these empirical DEGs were cross-referenced against a rigorously curated panel of canonical lineage markers compiled from associated literature and the CellMarker database [10–16]. The reference panel included markers for lymphoid lineages (B cells: *CD79A*; Plasma cells: *MZB1*; T cells: *CD3E*, *CD8A*, *CD8B*; NK cells: *XCL1*, *GNLY*) and myeloid cells (*CD163*). Finally, parenchymal and stromal populations were annotated utilizing canonical markers for hepatocytes (*ALB*, *APOA2*, *APOC3*, *CYP3A4*), cholangiocytes (*EPCAM*, *KRT19*), endothelial cells (*VWF*), and fibroblasts (*COL1A1*). Final cell-type identities were assigned based on a consensus between the automated CellTypist predictions and robust marker expression.

*Inference of Cell-Cell Communication Networks*

To systematically characterize intercellular communication across disease states, CellPhoneDB (v5.0) was employed via its Python pipeline for ligand–receptor (LR) interaction analysis[20]. The cpdb_statistical_analysis_method function was used to predict potential interaction strengths between cell types based on the expression levels of cognate LR pairs, retaining only ligand or receptor genes expressed in more than 10% of cells within a given type. Enriched interactions were determined through a permutation test in which cell type labels were randomly shuffled 1,000 times to compute null distributions, and significant LR pairs were extracted at a P value threshold of 0.05. For each disease, a consensus cell-cell interaction (CCI) network was derived by aggregating results across multiple subsampled iterations: a cell type pair was assigned an interaction edge only if at least one significant LR pair was detected (cutoff $\geq 1$) and the interaction was reproduced in $\geq$90% of iterations, thereby ensuring biological robustness.

*Construction of the Heterogeneous Meta-graph*

The disease-specific consensus CCI networks were integrated into a heterogeneous meta-graph comprising two node classes—disease nodes and cell type nodes—connected by two edge types. Cell-cell interaction edges were consolidated across diseases such that interactions shared by multiple conditions were annotated with the contributing disease list and weighted by the number of conditions in which they were observed. Disease-to-cell type edges were added on the basis of cellular composition within each disease category, retaining only cell types with $\geq$10 cells and excluding adjacent normal samples, with edge weights normalized to the maximum count. Pairwise Jaccard similarity indices were computed between disease-specific CCI networks to quantify the degree of shared versus unique intercellular communication programs across disease states; the resulting meta-graph was exported as node and edge tables for downstream network analysis.

#### LiverDCP: Multi-scale Contextualized Graph Learning

*Graph construction*

The metagraph (MG) $\mathcal{G}_{\mathrm{MG}} = (\mathcal{V}_{\mathrm{MG}}, \mathcal{E}_{\mathrm{MG}})$ is a directed heterogeneous graph. To represent the distinct biological roles of clinical conditions and cellular environments, we partition the nodes $\mathcal{V}_{\mathrm{MG}}$ into two typed sets:

- **Disease Nodes ($\mathcal{V}_D$):** Representing clinical conditions (e.g., HCC, MASH, Cirrhosis).
- **Cell-type Nodes ($\mathcal{V}_C$):** Representing immune and parenchymal cell populations.

The edge set $\mathcal{E}_{\mathrm{MG}}$ comprises $M$ distinct semantic relation types. We precompute a set of metapath adjacency matrices $A_{\mathrm{MG}} = \{A_{CC}, A_{DC}\}$, where $A_{CC}, A_{DC} \in \{0,1\}^{|\mathcal{V}_{\mathrm{MG}}| \times |\mathcal{V}_{\mathrm{MG}}|}$ encodes a specific relationship, including disease-specific cell-cell interaction (CCI) and disease-cell interaction(DCI). To facilitate global information flow, we precompute the undirected neighbor set for each disease node $v \in \mathcal{V}_D$:

$$\mathcal{N}_{\mathrm{diag}}(v_d) = \{u \in \mathcal{V}_{\mathrm{MG}} \mid (v,u) \in \mathcal{E}_{\mathrm{MG}}^{\mathrm{undirected}}\}$$

These neighbor sets drive the disease propagation mechanism within the graph fusion convolution layers, where disease-node embeddings are iteratively updated by averaging their neighbors' embeddings.

We define a context as a unique (disease, cell-type) pair. For each context, we construct a homogeneous protein-protein interaction (PPI) graph $\mathcal{G}_{\mathrm{k}} = (\mathcal{V}_{\mathrm{k}}, \mathcal{E}_{\mathrm{k}}), k = 1,2, \dots 286$. While the topology of each is homogeneous, the global heterogeneity is captured by the multiplicity of total contexts.

For each context $k$, we define a local-to-global mapping $\phi_k : \mathcal{V}_{\mathrm{k}} \to \mathcal{V}_{\mathrm{global}}$, where $\mathcal{V}_{\mathrm{global}}$ is the set of all unique proteins across all contexts. This allows us to construct the local adjacency matrix $A_{PPI}^{(k)} \in \{0,1\}^{|\mathcal{V}_{\mathrm{k}}| \times |\mathcal{V}_{\mathrm{k}}|}$ and retrieve features from a shared protein universe.

Before the start of convolution, we initialize the input feature matrices $\boldsymbol{X}_{MG}$ and $\boldsymbol{X}_{PPI}$ to serve as the initial hidden states $H_{MG}^{(0)}$and $H_{PPI}^{(0)}$.

For each protein $p \in \mathcal{V}_{\mathrm{global}}$, we sample a baseline feature vector $\mathbf{f}_p^{\mathrm{init}} \sim \mathcal{N}(\mathbf{0}, \mathbf{I}_{d_f})$, where $d_f = 1024$. The context-specific feature matrix $\boldsymbol{X}_{PPI}^{(k)}$ is then formed by collecting the vectors for all $p \in \mathcal{V}_k$.

The metagraph feature matrix $\boldsymbol{X}_{MG} \in \mathbb{R}^{|\mathcal{V}_{MG}| \times d_f}$ is initialized using cellular deconvolution proportions. For a disease node $d$, the feature vector contains the proportions of each cell type $c \in \mathcal{V}_C$ observed in that condition, padded with Gaussian noise $\eta \sim \mathcal{N}(0, 0.01)$ to match dimension $d_f$.

To bridge the local and global scales, we prepare a bipartite mapping index $\mathcal{M}_{P \to C}$. For every protein $i$ in a local context $k$, $\mathcal{M}_{P \to C}$ stores the index of the corresponding cell-type node $v \in \mathcal{V}_C$ in the global metagraph. This index enables the "Up" pooling of protein signals to the metagraph and the "Down" broadcasting of clinical context back to the proteins.

*Model convolution architecture*

The architecture consists of $L$ stacked blocks. Each block $l$ takes the previous hidden states $\boldsymbol{H}_{PPI}^{(l-1)}$ and $\boldsymbol{H}_{MG}^{(l-1)}$ and updates them using the precomputed topological structures. The graph fusion convolution(F-GCN) module ("Up") serves two purposes: aggregating multi-relational information within the metagraph and pulling molecular signals upward from the PPI networks. For each of the precomputed metapath adjacency matrix $A_{CC}$or $A_{DC}$ we compute a relation-specific representation $\mathbf{Z}_m$ using a GAT kernel:

$$\mathbf{Z}_{CC} = \text{Activation}\left(\text{GAT}\left(\boldsymbol{H}_{MG}^{(l-1)}, A_{CC}\right)\right)$$
$$\mathbf{Z}_{DC} = \text{Activation}\left(\text{GAT}\left(\boldsymbol{H}_{MG}^{(l-1)}, A_{DC}\right)\right)$$

The representations are then fused using the semantic attention mechanism. We learn an importance weight $\beta_m$ for each metapath to produce a unified metagraph feature:

$$\beta_m = S\text{oftmax}(\mathbf{q}^T \tanh(\mathbf{W}_{\text{sem}}\mathbf{Z}_m + \mathbf{b})), m = CC, DC$$
$$\boldsymbol{H}_{MG}^{fusion} = \beta_{CC} \cdot \mathbf{Z}_{CC} + \beta_{DC} \cdot \mathbf{Z}_{DC}$$

Where $\mathbf{b} \in \mathbb{R}^{d_{\text{sem}}}$, and $\mathbf{q} \in \mathbb{R}^{d_{\text{sem}}}$ are learnable semantic attention parameters shared across all metapaths.

To ground the metagraph in local molecular states, we use the bipartite mapping index $\mathcal{M}_{P\to C}$ to pool each protein $i$'s embeddings into their corresponding cell-type node $c$:

$$\boldsymbol{H}_{MG}^{(l)} = \text{Activation}\left(\boldsymbol{W}_{up}\left[\boldsymbol{H}_{MG}^{fusion} || \sum_{i\in\mathcal{V}_c} \alpha_i \cdot \boldsymbol{H}_{PPI}^{(l-1)}\right] + \boldsymbol{b}\right)$$

Where || denotes the concatenation of the global semantic context and the aggregated local molecular state. $\boldsymbol{W}_{up}$ is a learnable weight matrix that projects the concatenated features back into the model's hidden dimension.

Once the metagraph $\boldsymbol{H}_{MG}^{(l)}$ is updated with global disease context, the PPI Enhancement module ("Down") propagates this information back to the protein level to refine the interactome representations.

Using the same mapping index $\mathcal{M}_{P\to C}$, the updated cell-type and disease context is projected back to each protein node. We reuse the attention weights $\alpha_i$ from the Up phase to ensure a symmetric exchange of information:

$$\boldsymbol{H}_{PPI,i}^{context} = \alpha_i \cdot \boldsymbol{H}_{MG,c}^{(l)}$$

By using the exact same $\alpha_i$ cached from the Up phase, model ensures that "key" proteins (the hubs or markers of a cell type) receive a stronger signal from the global disease context.

Finally, each protein node updates its state by aggregating information from its immediate neighbors in the context-specific PPI adjacency matrix $A_{PPI}^{(k)}$, conditioned on the newly received global context:

$$\boldsymbol{H}_{PPI}^{(l)} = \text{GAT}\left(\boldsymbol{H}_{PPI}^{(l-1)} + \boldsymbol{H}_{PPI}^{context}, A_{PPI}^{(k)}\right)$$

Collectively, the complete forward pass for a single block can be summarized as a coupled update:

$$\left(\boldsymbol{H}_{PPI}^{(l)}, \boldsymbol{H}_{MG}^{(l)}\right) = \text{LiverDCP\_block}\left(\boldsymbol{H}_{PPI}^{(l-1)}, \boldsymbol{H}_{MG}^{(l-1)}, A_{PPI}, A_{CC}, A_{DC}, \mathcal{M}_{P\to C}\right)$$

*Embedding Quality Metrics*

To ensure that the learned embeddings capture biologically meaningful context-specific representations, we monitored five complementary embedding quality metrics at every training epoch. These metrics collectively assess whether distinct PPI contexts are mapped to well-separated, compact clusters in embedding space.

**Inter-Context Cosine Similarity.** We measured the distinguishability of context representations by computing the mean pairwise cosine similarity between context centroids. The centroid of context $k$ is $\bar{\boldsymbol{h}}_k = \frac{1}{N_k}\sum_{i=1}^{N_k} \boldsymbol{h}_{k,i}$. The metric is defined as:

$$\text{InterCtxSim} = \frac{1}{\binom{C}{2}} \sum_{i<j} \frac{\bar{\boldsymbol{h}}_i \cdot \bar{\boldsymbol{h}}_j}{\|\bar{\boldsymbol{h}}_i\| \, \|\bar{\boldsymbol{h}}_j\|}$$

Values near zero or negative indicate strong context separation, while values approaching 1 suggest embedding collapse.

**Silhouette Score.** For each protein embedding $\boldsymbol{x}_i$ with context label $\ell_i$, the silhouette coefficient is $s(i) = \frac{b(i)-a(i)}{\max\{a(i),b(i)\}}$, where $a(i)$ is the mean intra-cluster distance and $b(i)$ is the mean distance to the nearest foreign cluster. The overall silhouette score $\bar{s} = \frac{1}{N}\sum_i s\,(i)$ ranges from $-1$ to $1$, with higher values indicating denser, better-separated clusters.

**Calinski–Harabasz Index.** The variance ratio criterion quantifies the ratio of between-group to within-group dispersion:

$$\text{CH} = \frac{\text{Tr}(\boldsymbol{B})}{\text{Tr}(\boldsymbol{W})} \cdot \frac{N-C}{C-1}$$

where $\boldsymbol{B}$ and $\boldsymbol{W}$ are the between-group and within-group dispersion matrices, respectively. Higher values indicate better-defined clusters.

**Davies–Bouldin Index.** This index penalizes clusters that are internally spread out yet close to neighboring clusters:

$$\text{DB} = \frac{1}{C}\sum_{k=1}^{C} \max_{l \neq k} \frac{s_k + s_l}{d_{kl}}$$

where $s_k = \frac{1}{N_k}\sum_{i \in C_k} \|\boldsymbol{x}_i - \bar{\boldsymbol{h}}_k\|$ is the mean scatter of cluster $k$ and $d_{kl} = \|\bar{\boldsymbol{h}}_k - \bar{\boldsymbol{h}}_l\|$ is the inter-centroid distance. Lower values indicate more compact, well-separated clusters.

**Composite Embedding Quality Score.** We defined a composite quality score $Q \in [0,1]$ that aggregates the two most informative sub-metrics:

$$Q = \frac{1}{2}\left[(1 - \text{InterCtxSim}) + \frac{\bar{s}+1}{2}\right]$$

where the first term captures centroid-level separation and the second normalizes the silhouette score to $[0,1]$. This score served as the primary signal for triggering the Phase 1 to Phase 2 transition: when $Q$ exceeded a threshold $\tau_{\text{quality}}$ (default 0.7), the best-quality checkpoint was saved. The quality score was also monitored alongside ROC-AUC to detect regressions in cluster structure during link-prediction optimization.

*Training Protocol and Hyperparameters*

Optimization Strategy: The model was trained using the Adam optimizer with a base learning rate of $\eta = 1 \times 10^{-3}$ and weight decay of $5 \times 10^{-4}$. To stabilize the optimization of the non-convex center loss landscape, we employed a Dynamic Lambda Scheduling strategy. The regularization weight $\lambda_c$ was subjected to a linear warmup over the first $T_{\text{warmup}} = 15$ epochs, preventing strong clustering forces from distorting the initial topological learning.

Learning Rate Schedule: We utilized a cosine annealing schedule with warm restarts. The learning rate increases linearly during the warmup phase and then decays following a cosine curve:

$$\eta_t = \eta_{\text{min}} + \frac{1}{2}(\eta_{\text{max}} - \eta_{\text{min}})(1 + cos(\frac{t - T_{\text{warmup}}}{T_{\text{max}}}\pi))$$

Computational Implementation: Training was performed on a single NVIDIA GTX 5090 GPU using PyTorch Geometric. To handle the massive adjacency matrices of liver-scale interactomes ($> 10^5$ edges), we implemented a memory-efficient chunked inference algorithm for the bilinear link predictor, processing edges in batches of 4,096 to constrain peak memory usage.

### Supplementary Figures

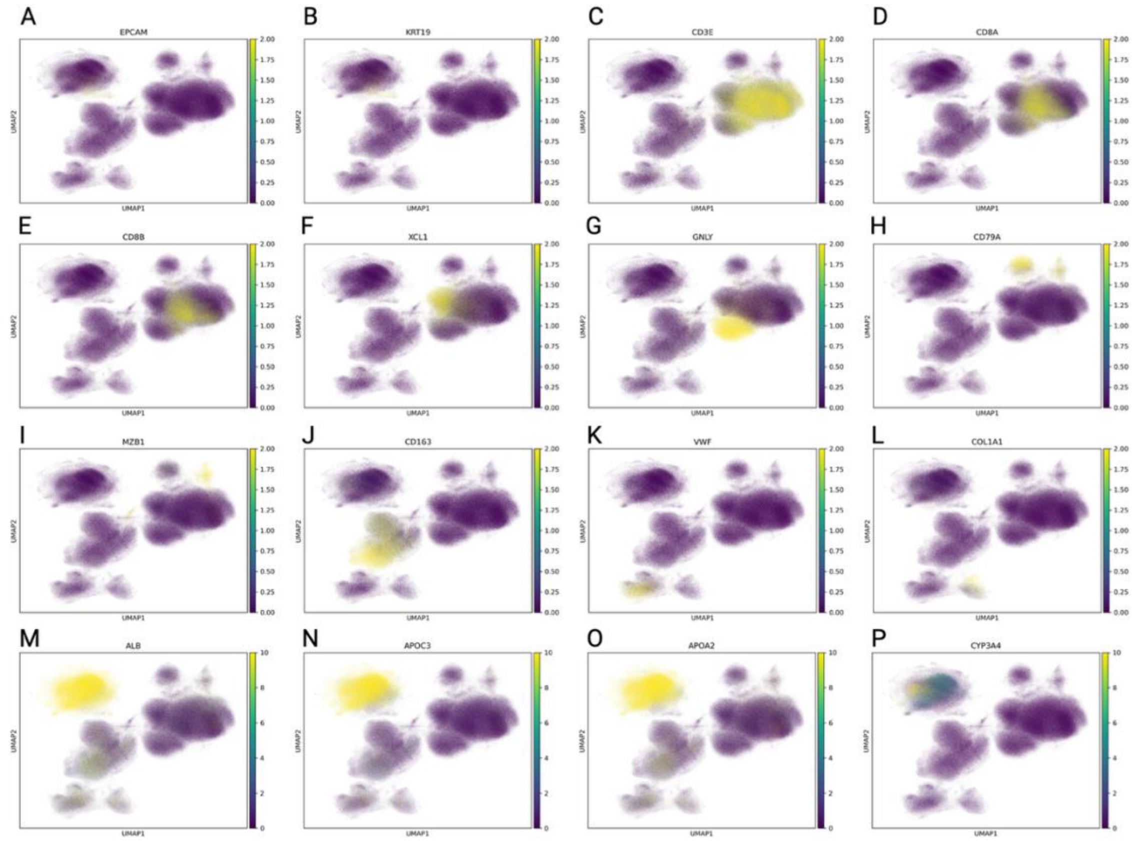


**Figure S1 | Canonical marker gene expression defining major cellular lineages across the LiverHomo atlas.**

**A–P**, Uniform Manifold Approximation and Projection (UMAP) embeddings of 1.16 million harmonized single cells from the LiverHomo integrated atlas. Cells are colored by the normalized transcript expression levels of established cell-type-specific marker genes utilized for unsupervised clustering and subsequent automated annotation. **A, B**, Cholangiocyte markers (*EPCAM*, *KRT19*). **C–G**, Lymphoid markers delineating T cells and natural killer (NK) cells, including pan-T (*CD3E*), cytotoxic CD8+ T cell coreceptors (*CD8A*, *CD8B*), and effector/cytotoxic molecules (*XCL1*, *GNLY*). **H, I**, B cell (*CD79A*) and plasma cell (*MZB1*) markers. **J**, Myeloid cell marker (*CD163*). **K**, Endothelial cell marker (*VWF*). **L**, Hepatic stellate cell (HSC) and mesenchymal marker (*COL1A1*). **M–P**, Hepatocyte markers reflecting core parenchymal synthetic and metabolic functions (*ALB*, *APOC3*, *APOA2*, *CYP3A4*). Color scales represent log-normalized gene expression.

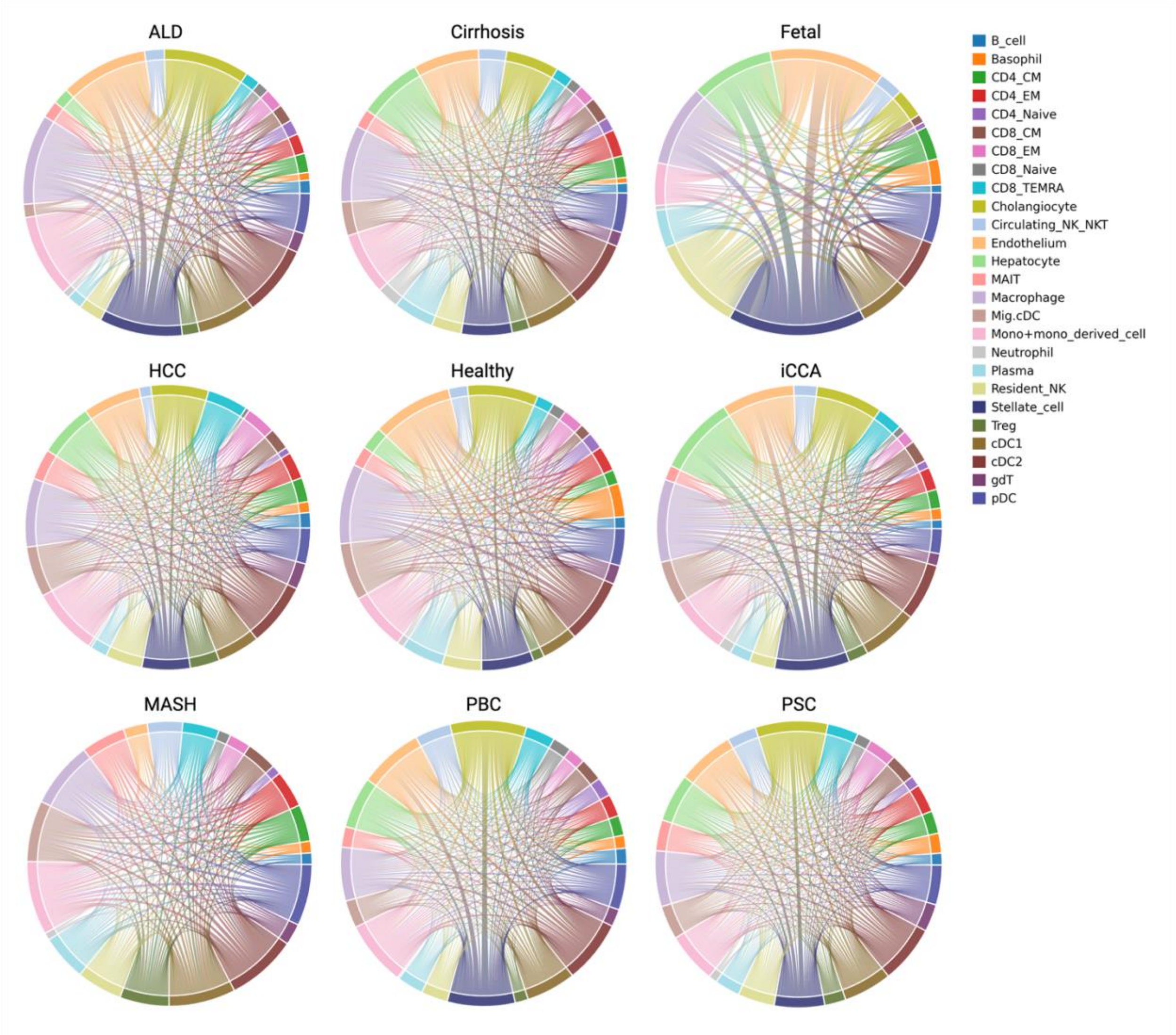


**Figure S2 | Global intercellular communication networks across human liver health, development, and disease.**

Chord diagrams illustrating the comprehensive cell-cell interaction (CCI) landscapes within healthy, developmental (Fetal), and distinct pathological liver environments (ALD, Cirrhosis, HCC, iCCA, MASH, PBC, and PSC). Intercellular signaling networks were inferred by integrating cell-type-specific transcriptomic profiles from the LiverHomo atlas with curated ligand–receptor pairs via CellPhoneDB. The outer peripheral ring segments correspond to 26 annotated high-resolution cell lineages and subtypes, color-coded according to the adjacent legend. The internal chords connect interacting cellular compartments, with the thickness of each connecting arc proportional to the total number of significant inferred ligand–receptor interactions between the corresponding cell populations. These global interactome models highlight the extensive topological rewiring of stromal, immune, and parenchymal cross-talk that drives disease progression and hepatic microenvironmental remodeling.

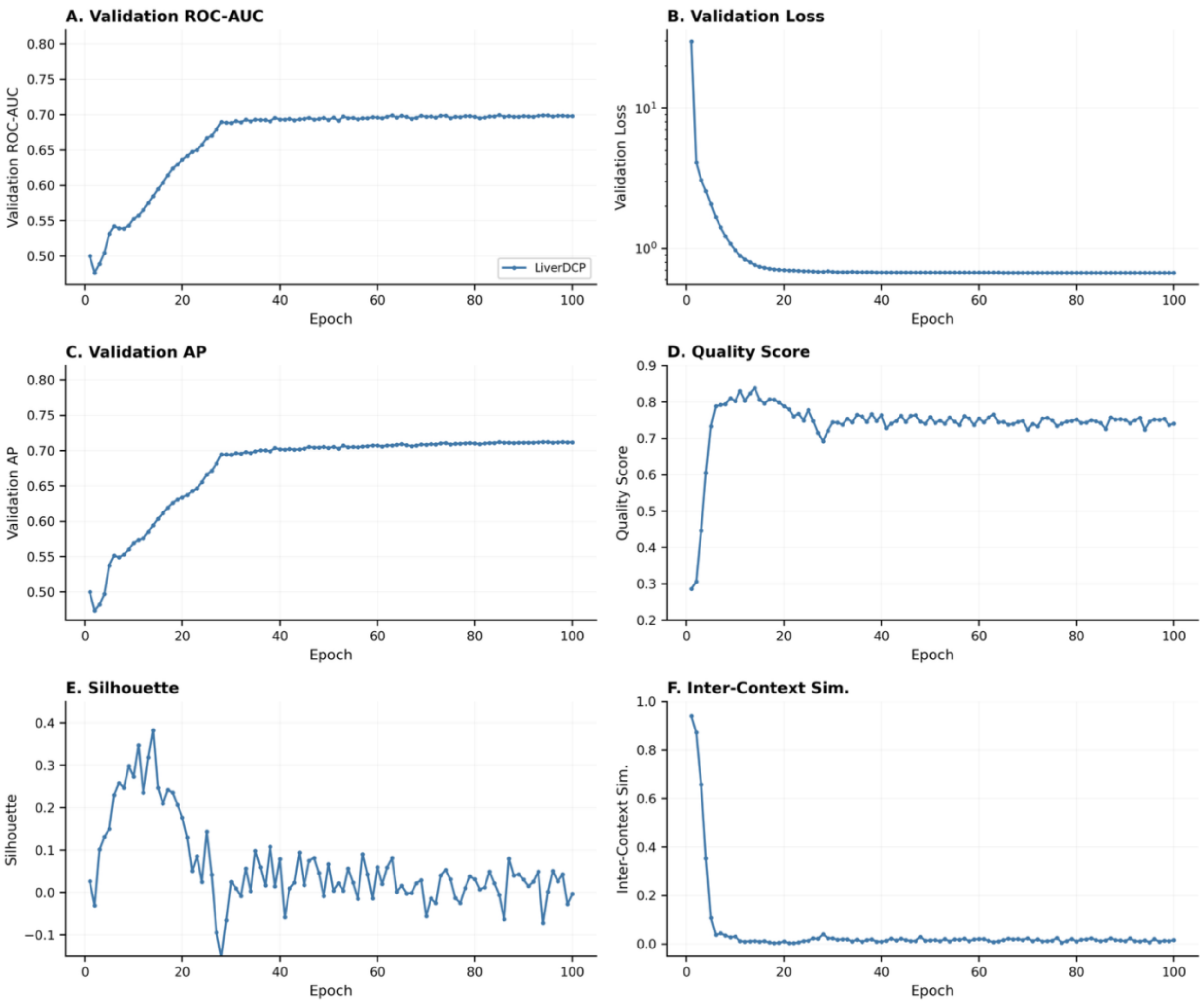


**Figure S3 | Training dynamics and optimization trajectories of the LiverDCP architectures.**

A & C) Evolution of predictive generalization, measured by validation ROC-AUC (A) and validation average precision (AP) (C). Both metrics exhibit a steady ascent during the initial 30 epochs before stabilizing near 0.70, demonstrating the model's robust capacity to learn and generalize topological link predictions. B) Validation Loss trajectory. Plotted on a logarithmic scale, the loss demonstrates rapid, stable convergence within the first 20 epochs, with no evidence of late-stage overfitting. D-F)Quantitative assessment of embedding space geometry and context separation. The overall Quality Score (D) rises sharply and maintains a high plateau (~0.75). The Silhouette score (E) peaks early (around epoch 15) before oscillating at a lower baseline, directly illustrating the timescale mismatch where geometric clustering resolves earlier than fine-grained topological edge optimization. Crucially, Inter-Context Similarity (F) drops precipitously to near zero within the first 10 epochs and remains suppressed, confirming the model rapidly and permanently learns to differentiate distinct cellular environments without suffering from context collapse.

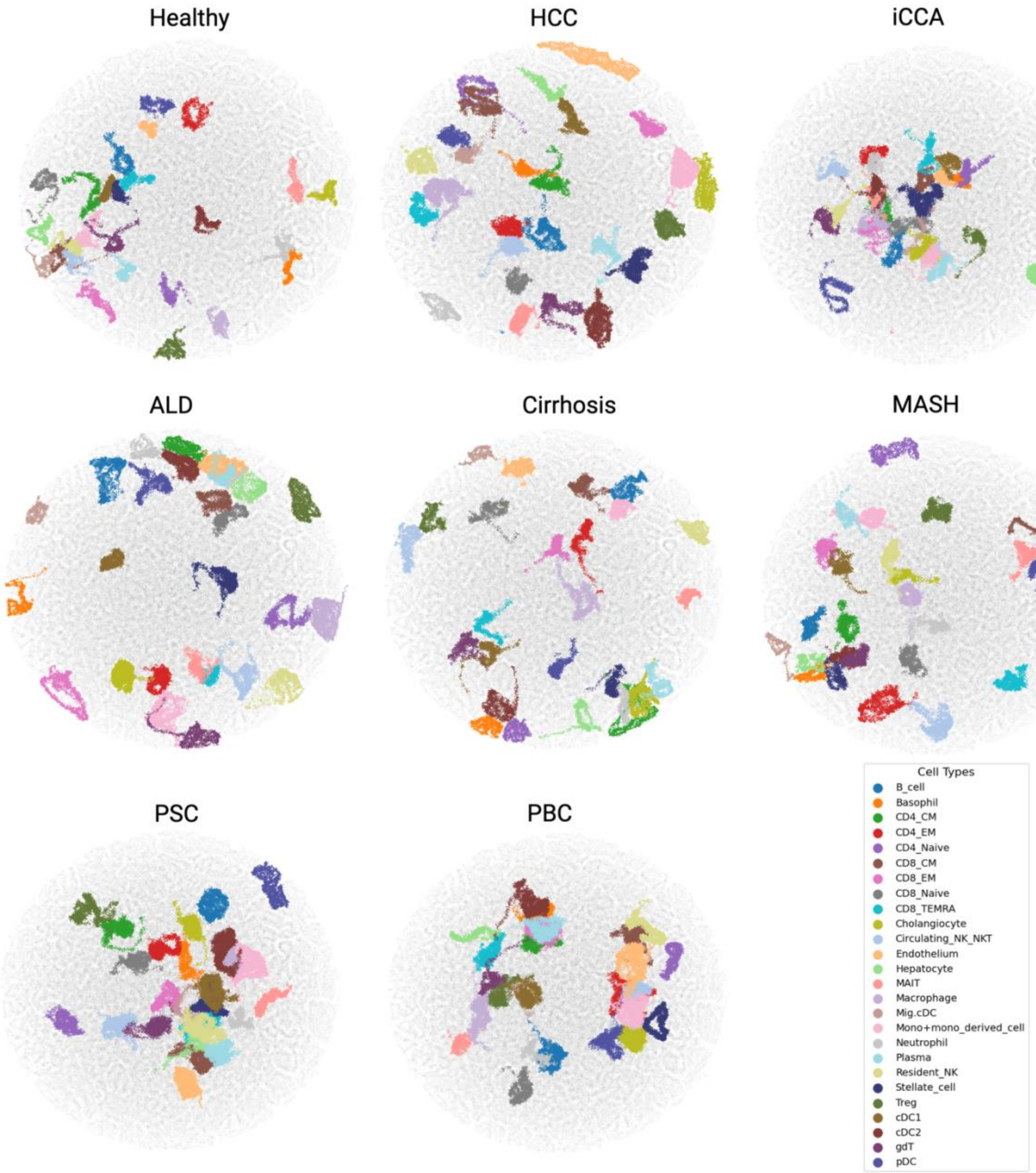


**Figure S4 | UMAP visualization of context-aware protein embeddings across distinct liver diseases.** UMAP of LiverDCP-generated protein representations, stratified by eight distinct clinical conditions: Healthy, HCC, iCCA, ALD, Cirrhosis, MASH, PSC, and PBC. Within each disease-specific manifold, protein embeddings are colored according to their corresponding cellular context (encompassing 26 distinct lineages, as indicated in the legend), with the gray background representing the global, non-target protein embedding space. The highly resolved, non-overlapping clustering of protein populations within each disease state demonstrates the efficacy of the model's geometric regularization constraints. This confirms that the architecture successfully deconvolutes the global interactome into highly specific cellular niches, maintaining robust intra-class compactness and inter-class separability across the diverse spectrum of hepatic health and disease.

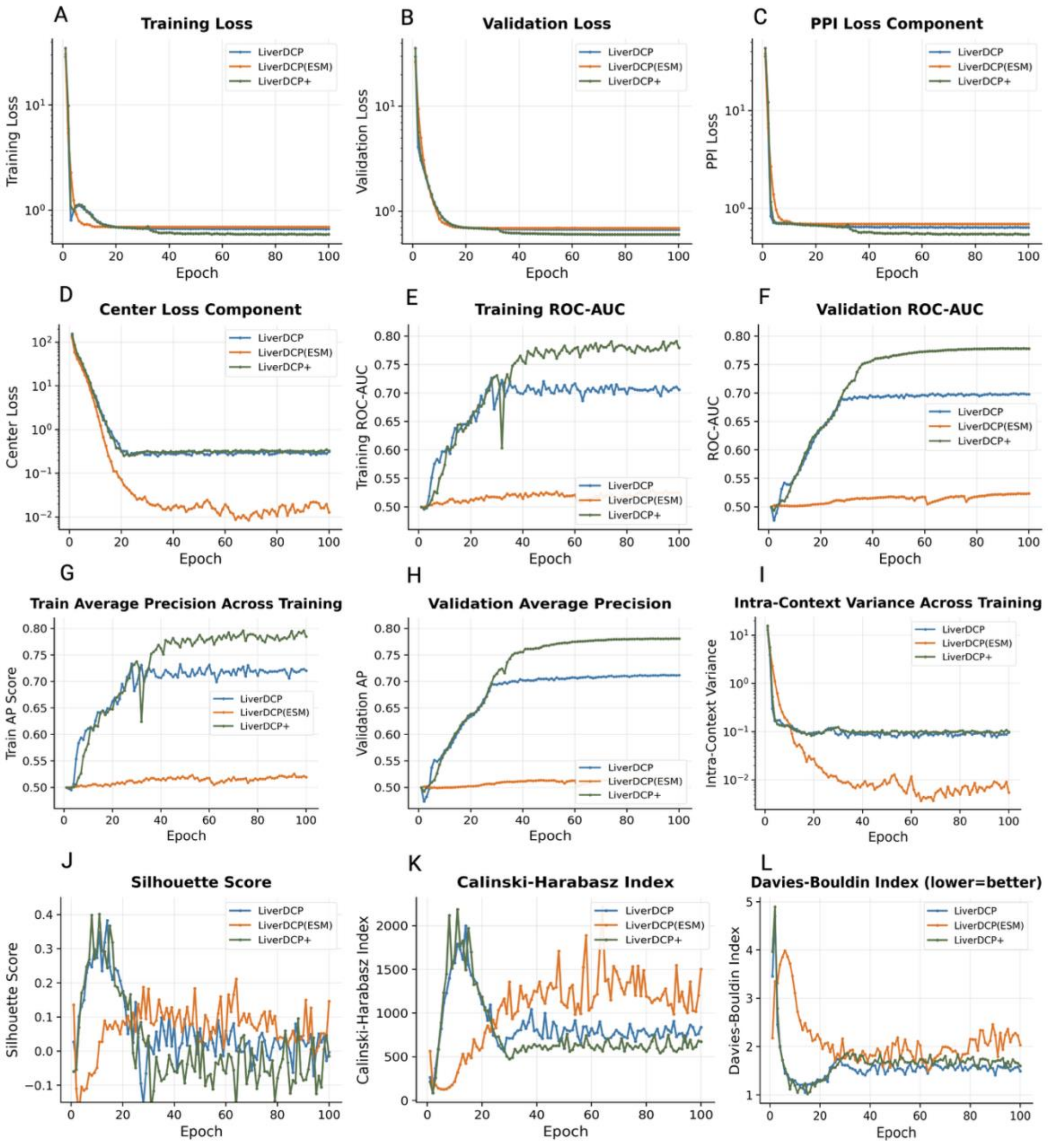


**Figure S5 | Training dynamics and optimization trajectories of the 3-version model.**

A & B) Evolution of the overall training loss (A), validation loss (B), and the specific protein-protein interaction (PPI) link prediction loss component (C) over 100 epochs. D & I) The comparison includes the topological baseline (LiverDCP, blue), the sequence-initialized model (LiverDCP(ESM), orange), and the hybrid flagship architecture (LiverDCP+, green). Trajectories of the center loss component (D) and intra-context variance (I). The sequence-only LiverDCP(ESM) model exhibits artificially low center loss and variance, which corresponds to "context collapse" (an inability to differentiate embeddings across distinct cellular niches). Conversely, LiverDCP+ balances variance to maintain cell-type specificity. E-H), Predictive performance trajectories measured by Training ROC-AUC (E), Validation ROC-AUC (F), Training Average Precision (G), and Validation Average Precision (H). LiverDCP+ progressively unlocks superior topological predictive power over the baseline, while the sequence-only model (orange) fails to capture the underlying interaction network. J–L) Quantitative assessment of latent space geometry over time using the Silhouette Score (J), Calinski-Harabasz Index (K), and Davies-Bouldin Index (L); lower values indicate superior clustering.

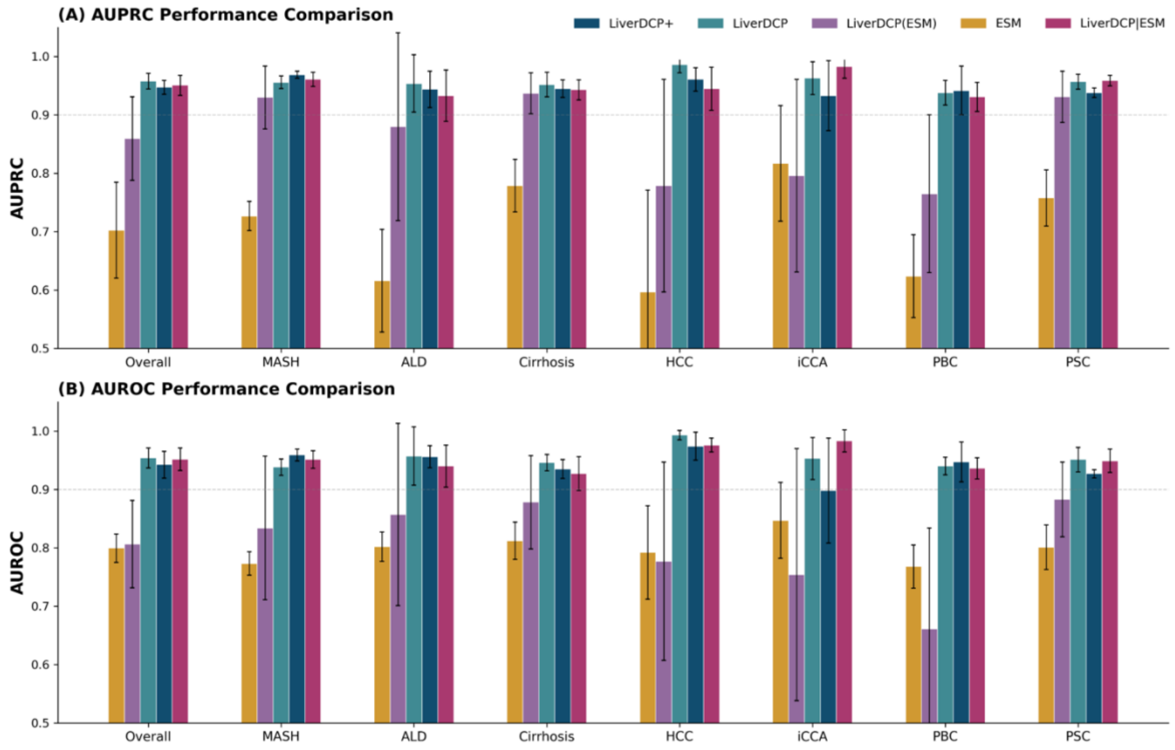


**Figure S6 | Superior prioritization of GWAS-derived disease risk genes using context-aware LiverDCP architectures.** Quantitative performance evaluation of five distinct modeling strategies for the binary classification of high-confidence GWAS risk loci against negative controls, measured by AUPRC (A) and AUROC (B). The 5-arm ablation study benchmarks the context-agnostic sequence baseline (ESM, gold) against four graph-based variants: a sequence-initialized graph model (LiverDCP(ESM), purple), the pure topological baseline (LiverDCP, teal), a static feature concatenation (LiverDCP|ESM, magenta), and the fully integrated hybrid flagship (LiverDCP+, dark blue). Across all seven diverse hepatic indications—spanning metabolic (MASH), autoimmune (PBC, PSC), and neoplastic (HCC) traits—architectures incorporating liver-specific graph priors (teal, dark blue, magenta) consistently and dramatically outperform the sequence-only baseline (gold). The pronounced performance gap and significantly reduced variance in the LiverDCP and LiverDCP+ models underscore a fundamental biological principle: genetic disease risk is heavily driven by the topology of the local interactome rather than intrinsic protein sequence alone. Error bars denote the standard deviation across 5 independent training seeds.

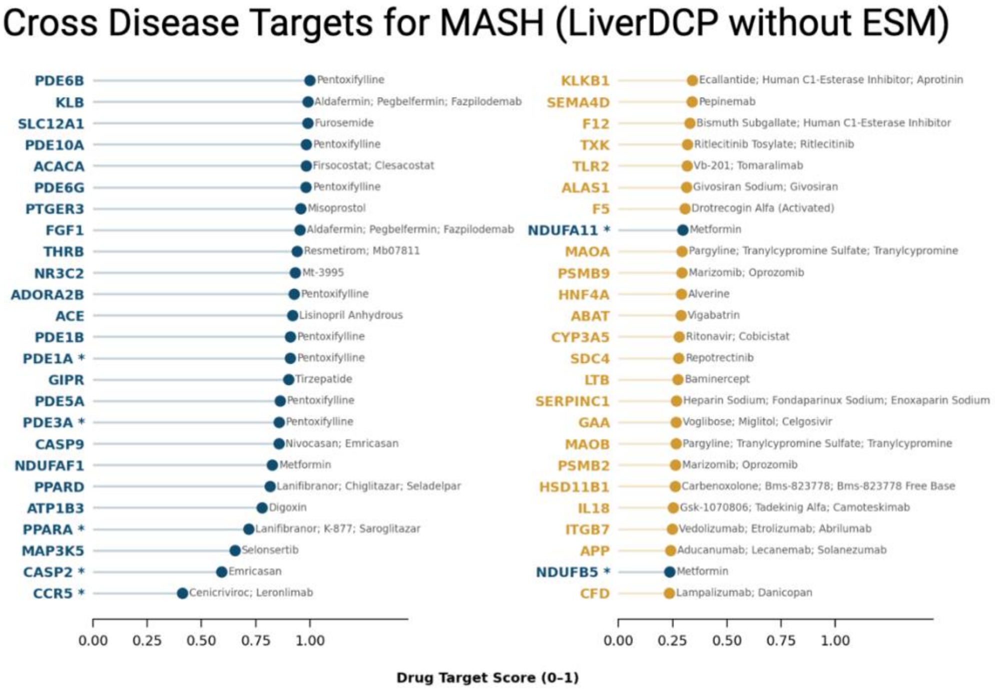


**Figure S7 | Cross-disease therapeutic target prioritization for MASH using the topology-only LiverDCP model.** Ranked prediction scores for the top 50 high-confidence candidate targets evaluated against the MASH drug-target centroid. This analysis utilizes the purely topological LiverDCP baseline embeddings—strictly excluding ESM-2 evolutionary sequence priors—to isolate the predictive power of the context-aware interaction network. Candidate proteins are stratified into known MASH Phase 2+ clinical targets (dark blue) and established pharmacological targets from other disease indications (gold) to highlight repurposing opportunities. The x-axis represents the normalized Mahalanobis proximity score (0–1). Even in the absence of sequence intelligence, the cell-type-specific graph topology successfully prioritizes highly relevant clinical targets. Notably, the model recovers THRB (the specific target of the first FDA-approved MASH therapeutic, resmetirom), alongside other advanced clinical targets such as PPARA (lanifibranor) and pan-caspase inhibitors (CASP2/9; emricasan), validating that disease-specific network wiring inherently captures critical pharmacological liabilities.

### Supplementary Tables

##### Table S1 | GWAS AUROC Performance for 5 Models

| Disease | LiverDCP+ | LiverDCP | LiverDCP(ESM) | ESM | LiverDCP\|ESM |
|---|---|---|---|---|---|
| MASH | 0.959 ± 0.010 | 0.938 ± 0.014 | 0.834 ± 0.123 | 0.773 ± 0.020 | 0.951 ± 0.015 |
| ALD | 0.956 ± 0.019 | 0.957 ± 0.050 | 0.857 ± 0.156 | 0.802 ± 0.025 | 0.940 ± 0.036 |
| Cirrhosis | 0.935 ± 0.016 | 0.946 ± 0.014 | 0.878 ± 0.080 | 0.812 ± 0.032 | 0.927 ± 0.029 |
| HCC | 0.974 ± 0.024 | 0.993 ± 0.008 | 0.777 ± 0.170 | 0.792 ± 0.080 | 0.976 ± 0.012 |
| iCCA | 0.898 ± 0.090 | 0.953 ± 0.036 | 0.754 ± 0.216 | 0.847 ± 0.065 | 0.983 ± 0.019 |
| PBC | 0.947 ± 0.034 | 0.940 ± 0.015 | 0.661 ± 0.173 | 0.768 ± 0.037 | 0.936 ± 0.018 |
| PSC | 0.927 ± 0.007 | 0.951 ± 0.021 | 0.883 ± 0.064 | 0.801 ± 0.038 | 0.949 ± 0.020 |

##### Table S2 | Drug 5-fold CV Performance

| Disease | Known | Train (90%) | LivePPI EF@10 | ESM EF@10 |
|---|---|---|---|---|
| HCC | 175 | 158 | 5.39 ± 0.72 | 4.97 ± 0.45 |
| MASH | 117 | 88 | 3.41 ± 0.43 | 2.73 ± 0.96 |
| iCCA | 73 | 66 | 4.44 ± 0.86 | 3.71 ± 0.49 |
| ALD | 29 | 27 | 3.25 ± 0.49 | 3.23 ± 0.35 |
| Cirrhosis | 59 | 54 | 4.63 ± 0.40 | 3.53 ± 0.70 |
| PBC | 37 | 34 | 4.06 ± 0.28 | 3.64 ± 0.52 |
| PSC | 18 | 17 | 2.24 ± 0.05 | 1.83 ± 0.36 |

**Table S3 | Top 50 Novel targets rank for MASH drug discovery**

| protein | score | category | rank |
|---|---|---|---|
| KAT6A | -0.9609699 | Novel | 1 |
| NIPA2 | -0.9959726 | Novel | 2 |
| PIK3CD | -0.99613756 | Novel | 3 |
| CLSTN1 | -0.9976533 | Novel | 4 |
| THOC3 | -0.9980495 | Novel | 5 |
| COPS3 | -0.9997659 | Novel | 6 |
| MOB3A | -1.0011178 | Novel | 7 |
| CDC40 | -1.009569 | Novel | 8 |
| CDC73 | -1.0097363 | Novel | 9 |
| PARL | -1.0099393 | Novel | 10 |
| SUDS3 | -1.0151485 | Novel | 11 |
| MAF | -1.0151802 | Novel | 12 |
| CUL4A | -1.0162569 | Novel | 13 |
| CARD11 | -1.0189315 | Novel | 14 |
| ITGB1BP1 | -1.0191061 | Novel | 15 |
| NARS1 | -1.0210893 | Novel | 16 |
| TET2 | -1.0223808 | Novel | 17 |
| ZNF148 | -1.0231556 | Novel | 18 |
| RIT1 | -1.025114 | Novel | 19 |
| EDEM1 | -1.0276276 | Novel | 20 |
| DDX3Y | -1.0297573 | Novel | 21 |
| HK1 | -1.030603 | Novel | 22 |
| SLC38A10 | -1.0344646 | Novel | 23 |
| SESN3 | -1.0345131 | Novel | 24 |
| GCLC | -1.0366257 | Novel | 25 |
| TGFBR1 | -1.0375562 | Novel | 26 |
| MYO9A | -1.0379642 | Novel | 27 |
| ATG5 | -1.0397104 | Novel | 28 |
| TMEM87A | -1.0423024 | Novel | 29 |

| | | | |
|---|---|---|---|
| GTF3C1 | -1.0424216 | Novel | 30 |
| MORC3 | -1.0426583 | Novel | 31 |
| DDHD1 | -1.0430257 | Novel | 32 |
| VAMP3 | -1.0445312 | Novel | 33 |
| LEO1 | -1.044774 | Novel | 34 |
| DYNC1LI1 | -1.0448602 | Novel | 35 |
| CTNNBL1 | -1.0454584 | Novel | 36 |
| NELL1 | -1.0463618 | Novel | 37 |
| CCNK | -1.0469637 | Novel | 38 |
| CHD8 | -1.0472802 | Novel | 39 |
| E2F4 | -1.0476089 | Novel | 40 |
| ZNF90 | -1.0493835 | Novel | 41 |
| THUMPD1 | -1.0498544 | Novel | 42 |
| PNP | -1.0502393 | Novel | 43 |
| ARHGEF2 | -1.0502517 | Novel | 44 |
| TAF11 | -1.0509275 | Novel | 45 |
| PFDN4 | -1.0509784 | Novel | 46 |
| COQ10B | -1.0514584 | Novel | 47 |
| CD9 | -1.0524504 | Novel | 48 |
| KPNA1 | -1.0531495 | Novel | 49 |
| OAT | -1.0539279 | Novel | 50 |
